# Spatial and temporal autocorrelation weave complexity in brain networks

**DOI:** 10.1101/2021.06.01.446561

**Authors:** Maxwell Shinn, Amber Hu, Laurel Turner, Stephanie Noble, Katrin H. Preller, Jie Lisa Ji, Flora Moujaes, Sophie Achard, Dustin Scheinost, R. Todd Constable, John H. Krystal, Franz X. Vollenweider, Daeyeol Lee, Alan Anticevic, Edward T. Bullmore, John D. Murray

## Abstract

High-throughput experimental methods in neuroscience have led to an explosion of techniques for measuring complex interactions and multi-dimensional patterns. However, whether sophisticated measures of emergent phenomena can be traced back to simpler low-dimensional statistics is largely unknown. To explore this question, we examine resting state fMRI (rs-fMRI) data using complex topology measures from network neuroscience. We show that spatial and temporal autocorrelation are reliable statistics which explain numerous measures of network topology. Surrogate timeseries with subject-matched spatial and temporal autocorrelation capture nearly all reliable individual and regional variation in these topology measures. Network topology changes during aging are driven by spatial autocorrelation, and multiple serotonergic drugs causally induce the same topographic change in temporal autocorrelation. This reductionistic interpretation of widely-used complexity measures may help link them to neurobiology.

## Introduction

The amount of data we can collect about the brain is growing rapidly, expanding the types of neuroscience that can be done. Until recently, the statistical methods used within neuroscience have focused on testing specific well-defined hypotheses on datasets collected specifically to test them. Now, new high-throughput methods for data collection, as well as the availability of large open-access datasets, provide exciting opportunities to understand how brain regions work together. We can now look for patterns in the data which were not specified *a priori*, as well as regularities which cannot be understood by looking at one neuron, circuit, region by itself.

For example, in resting-state fMRI (rs-fMRI), brain activity is recorded when subjects are not engaged in any task, with the aim of finding multi-dimensional patterns of the brain at rest. Much rs-fMRI methodology is based on functional connectivity (FC), the matrix of pairwise correlations across brain regions ^1^. Due to the successes in understanding the brain using FC ^2^, nonlinear methods have been developed to dig deeper into the FC matrix, examining questions such as complexity and emergent phenomena which can only be understood at higher levels of abstraction.

Network neuroscience is a popular abstract framework for understanding complexity within the brain, and can be used to find and quantify sophisticated and interpretable patterns across the entire FC matrix ^3–5^. In this framework, FC is interpreted as a network using graph theory, where nodes represent brain regions and edges represent the correlation in activity between those regions over time. These graphs are often binarized to include only the strongest edges, forming an unweighted, undirected graph. Network analysis enables the study of brain topology at a higher level of abstraction, with the ability to look at properties such as clustering, modular organization, and regional influence of brain-wide activity, quantified as statistics called “graph metrics” ^3,6^. This powerful framework for understanding brain network topology has profoundly influenced how we understand the organization of rs-fMRI activity ^3–5^. However, because several levels of graph theoretic abstraction are necessary to understand these phenomena, it can be difficult to determine what lower-level factors may have caused a change in the network’s topology. This makes it especially difficult to find a biological source for any trends observed at the network level. Thus, there is great interest in linking these network topology metrics to lower levels of abstraction, such as simple timeseries statistics.

Here, we show that two simple timeseries statistics, spatial autocorrelation (SA) and temporal autocorrelation (TA), explain a large fraction of individual variation in network topology. SA and TA are attractive features because they reflect a myriad of physical and biological mechanisms within the brain, including molecular ^7,8^, circuit ^9–11^, activity state ^12,13^ and organizational ^14–17^ properties. SA and TA are reliable across subjects and brain regions.

Next, we show that SA and TA are sufficient to explain graph metrics. We developed a spatiotemporal model of surrogate timeseries which preserves only SA and TA. After fitting the model, we found these features alone are sufficient to reproduce measures of network topology. This model can also be used to understand which spatiotemporal features are most important for driving changes in graph metrics due to healthy aging.

Finally, we show that SA and TA are biologically relevant and meaningful beyond their connection to graph topology. SA and TA correlate with subclinical symptoms of dementia. Causal pharmacological manipulations with different serotonergic psychedelics show the same topographic patterns of decrease in TA, which graph metrics do not capture. Overall, SA and TA are closely tied to complex network topology and are important properties of rs-fMRI.

## Results

### Low dimensionality of graph topology measures

We found that many measures of graph topology are correlated with each other across subjects in networks constructed from rs-fMRI. We analyzed multiple neuroimaging datasets utilizing diverse methodologies implemented by different teams, focusing on the Human Connectome Project (HCP) dataset ^18^, with validation in the Yale Test-Retest (TRT) ^19^ and Cambridge Centre for Ageing and Neuroscience (Cam-CAN) ^20^ datasets. In line with previous work, networks were constructed by thresholding the FC matrix to maintain only the 10% strongest connections and ensuring that no nodes are disconnected from the rest of the network. We used several graph metrics to quantify different aspects of network topology. This included the propensity for network hubs to connect to each other (assortativity), the average distance between nodes (global efficiency), the number of triangles (transitivity), and the propensity to organize into local communities within the broader network (modularity). We also considered two nodal metrics commonly averaged across nodes for each network: the number of triangles surrounding a node (clustering coefficient), and the mean path length for a node’s neighbors (local efficiency). Lastly, we considered two nodal graph metrics which were not averaged across nodes: the total number of connections a node makes (degree), and the number of paths in the network which pass through a given node (centrality).

In all datasets, graph metrics are highly correlated with each other across subjects (Figure 1b, S1a,i,q) ^21–24^. However, unweighted graph metrics, derived from binarized networks, are also highly correlated with the mean (mean-FC), variance (var-FC), and kurtosis (kurt-FC) of FC (Figure 1b, S1a,i,q). This is surprising because the FC binarization procedure destroys all explicit information about mean-FC, var-FC, and kurt-FC (see Methods for proof). Thus, some unobserved underlying factors must influence both the statistical moments of FC and the topology of the unweighted graph. In what follows, we demonstrate that spatial and temporal autocorrelation are two such factors.

**Figure 1.**
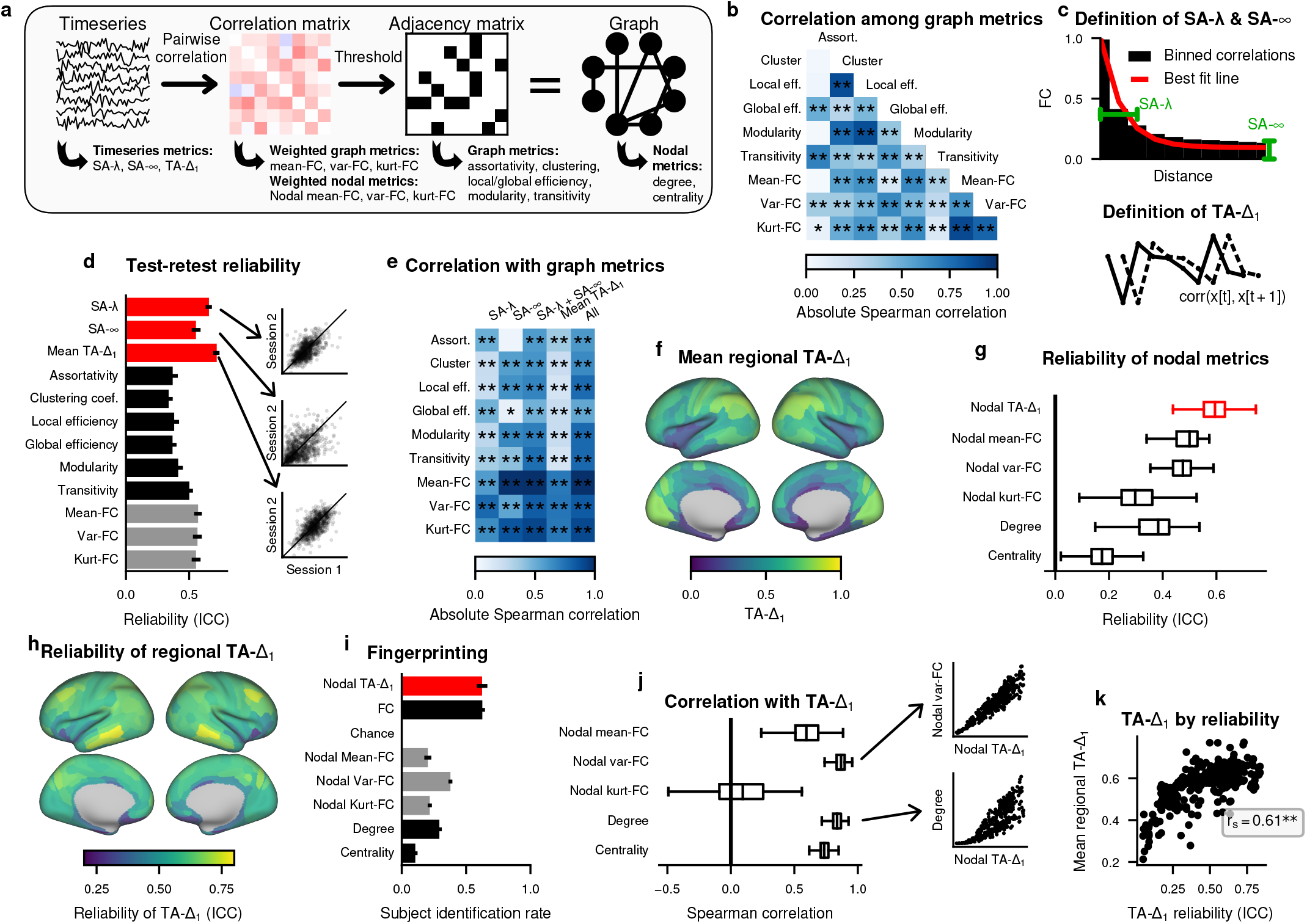
Spatial and temporal autocorrelation are important features of rs-fMRI timeseries. (a) Diagram describing the connectomic pipeline and key graph metrics. (b) Correlation of graph metrics across all subjects in the HCP dataset, N=883. * indicates p<0.05 and ** indicates p<0.01. (c) Schematic demonstrating the calculation of SA-λ and SA-∞ (top) and TA-Δ_1_ (bottom). (d) Test-retest reliability of graph metrics, quantified by intraclass correlation coefficient (ICC). Error bars indicate 95% confidence interval. Inset scatterplots show the correlation across subjects for two example sessions. (e) Correlation across subjects between weighted or unweighted graph metrics and SA-λ, SA-∞, or TA-Δ_1_. “SA-λ + SA-∞ “indicates a cross-validated linear model with these two terms, and “all” indicates a cross-validated linear model with all three terms. * indicates p<0.05 and ** indicates p<0.01. (f) Mean TA-Δ_1_ for each brain region. (g) Reliability is computed for each nodal graph metric separately in each region. Boxplots indicate the distribution of ICC across regions. Outliers hidden for visualization. (h) Reliability of TA-Δ_1_ is plotted across brain regions. (i) A fingerprinting analysis to identify individual subjects is computed for each of six possible pairs of the four sessions. The mean fraction of subjects correctly identified is shown, with error bars indicating standard error. (j) The correlation across regions of regional TA-Δ_1_ with nodal weighted and unweighted graph metrics was computed for each subject, and boxplots indicate the distribution across subjects for each metric. Outliers hidden for visualization. (k) TA-Δ_1_ for each region, shown in (f), is plotted against the reliability of TA-Δ_1_ for each region, shown in (h).

### Spatial autocorrelation

Spatial autocorrelation (SA) is the ubiquitous phenomenon in neuroscience that nearby regions are more similar than distant regions ^8,15,25–28^. In order to examine SA’s test-retest reliability, i.e. how well SA is preserved across different rs-fMRI sessions from the same subject, we developed a method to estimate SA on a single-subject level by decomposing it into two components: the rate at which FC falls off with physical distance (SA-λ), and the average correlation between two distant brain regions (SA-∞). Our method bins FC by distance and finds the best fit SA-λ and SA-∞ for each subject’s FC vs distance curve (Figure 1c), using Euclidean distance to accommodate both hemispheres and subcortex (Equation 1). Test-retest reliability is quantified with intraclass correlation coefficient (ICC), which compares the variability of multiple observations of the same subject to the variation across all subjects, where 1 is perfect reliability and 0 is chance.

We found that SA is both reliable and correlated with graph metrics across subjects. Both SA-λ and SA-∞ have high test-retest reliability compared to graph metrics (Figure 1d, S1b,j). The effect cannot be explained by head motion within the scanner (accounting for motion, partial correlation > .65, *p* < 10^−10^ for both SA-λ and SA-∞) or global signal regression (Figure S1), and is strongest in the HCP dataset, which had the smallest motion and parcel size confounds (Figure S3a,b). Both SA-λ and SA-∞ are highly correlated with weighted and unweighted graph metrics (Figure 1e, S1c,k,r). To test whether the combined influence of SA-λ and SA-∞ is greater than either one alone, we constructed a linear model including SA-λ, SA-∞, and a constant term, training it on 50% of subjects selected randomly to fit each graph metric. This model significantly predicts graph metrics on the held-out data (Figure 1e, S1c,k,r). Therefore, SA is reliable and topologically informative.

### Temporal autocorrelation

Temporal autocorrelation (TA) describes the smoothness, or memory, of the rs-fMRI timeseries over time, and is known to vary heterogeneously across brain regions ^9,12,14,17,29,30^ and influence FC ^13,31^ and graph topology ^32–34^. We quantify TA for each parcellated region in each subject as the Pearson correlation between adjacent time points of the region’s timeseries, i.e. the lag-1 temporal autocorrelation (TA-Δ_1_) (Figure 1c). TA-Δ_1_ is a simple non-parametric measure of TA, which is related to other TA measures such as parametric estimates of long-memory dynamics (Figure S2) (Supplement 1) ^30,35^. We observed highest TA-Δ_1_ in occipital and parietal regions, and lowest in limbic regions (Figure 1f, S1d,l,s).

TA is reliable across subjects, both at the whole brain and regional level. At the whole brain level, we computed each subject’s mean TA-Δ_1_ by averaging across all regions, and found that mean TA-Δ_1_ is highly reliable compared to graph metrics (Figure 1d, S1b,j). For each region, we also computed the reliability across subjects, and found that median regional reliability was higher for TA-Δ_1_ than for other nodal graph metrics (Figure 1g, S1e,m). The regional reliability of TA-Δ_1_ varies heterogeneously across the brain (Figure 1h) ^36^.

This reliability and heterogeneity suggests that TA-Δ_1_ could be used to identify individual subjects across the population. To identify subjects, we utilized a “finger-printing” analysis ^19,37^. For a given measure, such as TA-Δ_1_, we matched each session of each subject to the session with the highest Pearson correlation in that measure across regions, selecting among all sessions from all subjects. Then, we counted the number of pairs for which both sessions belonged to the same subject. We found that fingerprinting using regional TA-Δ_1_ identified the single matching session from the pool of 1765 sessions in over 62.4% of subjects, compared to 62.6% with a more traditional fingerprinting analysis using FC (Figure 1i), with similar success in other datasets (Figure S1f,n). We compared this to fingerprinting using several nodal graph metrics, which were unable to match the performance (Figure 1i, S1f,n). This indicates that TA-Δ_1_ is reliable enough to identify an individual subject from a population.

TA-Δ_1_ is highly correlated with graph topology at the individual and regional levels. At the individual level, we found a strong correlation between mean TA-Δ_1_ and graph metrics (Figure 1e). To test its influence in conjunction with SA, we developed a linear model incorporating TA-Δ_1_, SA-λ, SA-∞, and a constant, training it on 50% of randomly-selected subjects to predict each graph metric. This model significantly predicted all graph metrics (Figure 1e), and almost all graph metrics in the other datasets (Figure S1c,k,r). At the regional level, for each subject, we computed the Spearman correlation between regional TA-Δ_1_ and various graph metrics. We found that both weighted and unweighted nodal graph metrics were highly correlated with TA-Δ_1_ (Figure 1j, S1g,o,t). Most notably, a node’s degree, or the total number of connections a node makes to other regions, was predicted by TA-Δ_1_ with a median correlation of 0.89 (Figure 1j), and this relationship was not driven by parcel size (median partial correlation 0.83). Remarkably, this implies that network hubs can be discovered without examining the topology of network ^33,34^.

### Multiple sources of temporal autocorrelation

TA in fMRI can be shaped by multiple underlying factors, such as the noise level of a parcel. For any temporally-autocorrelated timeseries, adding uncorrelated noise reduces the TA. Therefore, if the entire brain had a slow underlying timescale, and some regions were noisier than others, then TA would be lowest in regions with the most noise. Thus, a region’s TA partially reflects the amount of noise present ^38^. Consistent with this hypothesis, we found that regions with the lowest TA-Δ_1_ show the lowest reliability in TA-Δ_1_ (Figure 1k, S1h,p). By this logic, TA may influence FC by altering the fraction of shared variance between pairs of regions ^39^. In what follows, we utilized the property that noise influences TA to build a spatiotemporal model for generating surrogate timeseries with regionally heterogeneous noise.

### Spatiotemporal model for surrogate timeseries

SA and TA are sufficient to replicate individual variation in network topology. We fit individual subjects with a parsimonious spatiotemporal surrogate timeseries model. The model operates at the level of the parcellated timeseries, meaning that comparisons may be drawn at multiple levels of the analysis pipeline (Figure 1a). Our model can be summarized in a small number of steps (Figure 2a). We generate long-memory (1 / *f* ^α^ spectrum) timeseries which have uniformly high TA. Concurrently, we spatially embed these timeseries according to brain geometry, introducing SA by increasing the correlation of nearby regions, while preserving the frequency spectra. This is possible using our newly-derived generalization of phase randomization (Supplement 2.1). Then, we add uncorrelated white noise with a region-specific variance, thereby lowering TA heterogeneously (Supplement 2.3). The resulting timeseries can then be analyzed the same way as rs-fMRI timeseries. Thus, in our model, all variation in graph topology must be caused by variation in SA and/or TA.

**Figure 2.**
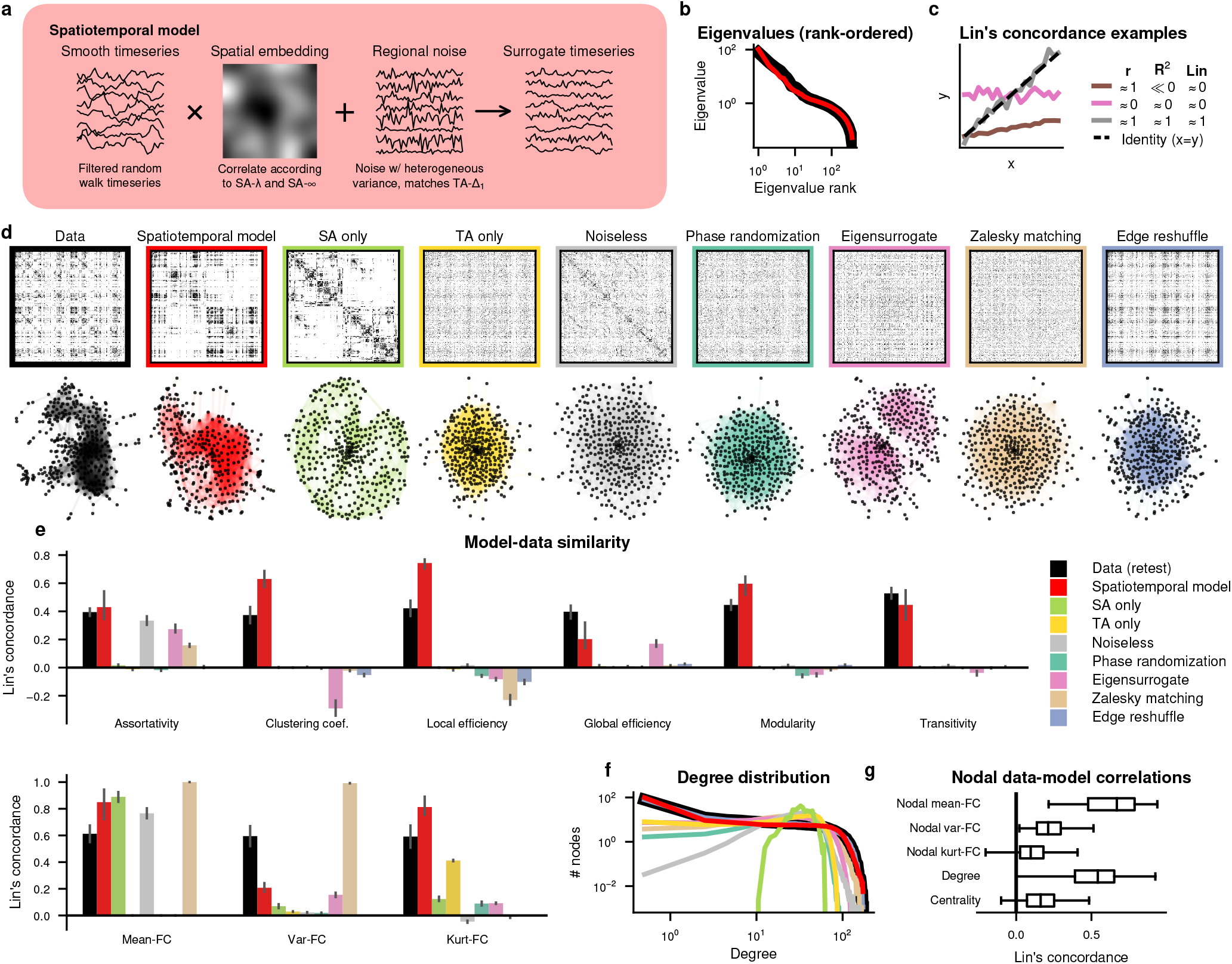
A spatiotemporal model captures connectome topology. (a) Schematic of our spatiotemporal modeling framework for surrogate timeseries with SA and regionally heterogeneous TA. (b) Log-log distribution of the eigenvalues of the FC matrix for an example subject (black) compared to the spatiotemporal model (red). (c) A schematic demonstrating Lin’s concordance, our model fit statistic. (d) Example FC matrices (top) and graphs (bottom) for the original data (black) and for each model (colors). Graphs are visualized using a force-directed layout, which positions topographically neighboring nodes nearby. (e) Lin’s concordance between model and data for each model. Bars represent the mean across the four scanning sessions, and error bars the standard error. For comparison, black indicates Lin’s concordance between sessions from the same subject. (f) Log-log degree distribution for each model compared to the data (black). (g) Distribution of Lin’s concordance of nodal metrics between model and data for each region. Outliers hidden for visualization.

Our model contains two SA parameters which are fit on the single-subject level, corresponding to noiseless SA-λ (SA-λ^gen^) and noiseless SA-∞ (SA-∞^gen^). We also utilize the subject’s observed regional TA-Δ_1_ to determine the variance of noise to add. Since it was not possible to estimate SA-λ^gen^ and SA-∞^gen^ directly from the data (Supplement 2.4), we fit them through optimization to a feature of the data we do not directly assess, the eigenvalue distribution, which is well-captured by the model (Figure 2b). This parameterization ensures that all variation in graph topology must be caused by variation in SA and/or TA.

### Model performance

We evaluated the ability of our model to capture weighted and unweighted graph metrics using surrogate timeseries. In principle, model fit could be assessed using several different criteria, such as the model’s ability to capture as much variance as possible (e.g. using *R*^2^), or alternatively, by its ability to capture individual variation (e.g. using Pearson correlation) (Table S1). Here, we assess model fit using Lin’s concordance, a stricter criterion which is 1 only when the model captures both variance and individual variation, and zero or negative when either assessment is poor (Figure 2c). An example subject’s FC matrix and network show qualitative similarities to the spatiotemporal model’s surrogate FC matrix and network Figure 2d.

Our model captures important features of graph topology. As an estimate of the total reliable variability of the measure, we compare the model to the Lin’s concordance of the measure on two independent sessions from the same subject. As expected, we confirmed the model matches the SA and TA of the original subjects (Figure S5). The model captures weighted and unweighted graph metrics, with similar Lin’s concordance as a second session from the same subject (Figure 2e, S6). It also reproduces the degree distribution (Figure 2f), i.e., the fraction of nodes with a given degree, and captures some regional graph metrics (Figure 2g).

To confirm that a simpler model is unable to match this performance, we fit versions of the model which capture only SA (“SA only”), only TA (“TA only”), or both SA and TA but with TA determined by intrinsic timescale instead of noise (“Noiseless”) (Figure S4). We also fit several commonly used null models: one which preserves the power spectrum amplitudes (“Phase randomization”), one which preserves mean-FC and var-FC (“Zalesky matching”) ^32^, and one which preserves degree distribution (“Edge reshuffle”) (Figure S4). Example surrogate FC matrices and networks on the same subject for these models show less qualitative similarity to the subject’s FC than the spatiotemporal model (Figure 2d). Likewise, none of these alternative models capture network topology as measured by weighted and unweighted graph metrics (Figure 2e, S5, S6, Table S1) or the degree distribution (Figure 2f). The ability of our spatiotemporal model to generally fit the Yale-TRT and Cam-CAN datasets better than alternative models underscores its generality (Figure S7a,d,g, Table S1).

To confirm whether our excellent fit was driven by our use of eigenvalues in the loss function, we created an “eigensurrogate” method for creating surrogate FC matrices. Eigensurrogates preserve the eigenvalue distribution of the FC matrix but randomize the eigenvectors, thereby perfectly duplicating the linear dimensionality of the FC matrix. This tests the null hypothesis that an effect is due to restrictions in dimensionality caused by SA and TA. We found that eigensurrogates were unable to reproduce the graph topology (Figure 2e, S5, S6, Table S1). They were also unable to reproduce graph properties in the HCP-GSR and Cam-CAN datasets (Figure S7a,g), but showed substantially more explanatory power than all other models in the Yale-TRT dataset (Figure S7d).

### Linking autocorrelation to graph topology

Our results show that SA and TA predict graph topology, but establishing this relationship mathematically is difficult due to the nonlinearity of constructing and analyzing a thresholded graph. Instead, we attempt to provide an intuition of why SA and TA may influence topology by considering their impact on individual edges. The impact of SA is relatively straightforward—SA increases the mean correlation between nearby regions, and thus, increases the probability of an edge. However, it is not immediately clear how TA might influence topology.

We examine here a statistical explanation for why strong TA may lead to high degree, consistent with our (Figure 1j) and others’^33,34^ previous finding that a node’s degree is highly correlated with TA-Δ_1_. Degree is determined by the number of correlations the node makes which exceed the binarization threshold (Figure 1a). This means that even if the expected value of the correlation (nodal mean-FC) is low, there may still be several correlations above the threshold if it has a high variance (nodal var-FC). In other words, a high variance may increase the probability of crossing the threshold moreso than a high mean. This means that any process which increases a node’s variance should also increase a node’s degree. Two temporally autocorrelated time-series are expected to have a higher variance in their pair-wise correlation ^34^ (Supplement 3), and this relationship is reflected in our data (Figure 1e,j). Thus, for individual nodes, one way that TA drives a node’s degree is by increasing the var-FC. If region A is highly correlated with regions B and C, it is also more likely that B and C are correlated with each other ^32^. This means that high– TA-Δ_1_ nodes in the graph are more likely to share an edge ^34,40^, creating a clustered network topology.

We confirmed this reasoning using a two-parameter “economical clustering” model ^22^, which builds graphs directly by probabilistically connecting nodes based on their distance and clustering topology (Supplement 4). This model is known to reproduce several topological features of brain networks ^22–24^, and is convenient because the procedure for constructing networks is distinct from the rs-fMRI pipeline. We found that changes in SA and TA correspond to an increased propensity for short and clustered edges, respectively, in the economical clustering model (Supplement 4) (Figure S8). The impact of these parameters on graph metrics is also qualitatively similar in the two models (Figure S8). In other words, the parameters of a graph-level generative process known to reproduce topological features of rs-fMRI networks are closely linked to the dimensions of network topology spanned by SA and TA. Thus, SA and TA can be interpreted directly in terms of graph topology.

### Spatial autocorrelation in healthy aging

We have shown that our surrogate model can trace differences in graph metrics among subjects down to differences in the SA and TA of the timeseries used to produce them. Therefore, we ask if the opposite is possible: if graph metrics change across a population, is SA or TA responsible for those changes in graph metrics? To do this, we look at age-related changes in graph metrics, and show that perturbations in the SA-∞ parameter, but not the SA-λ or TA-Δ_1_ parameters, lead to a similar change in graph metrics.

We analyzed the Cam-CAN dataset, containing crosssectional rs-fMRI data from over 800 subjects ranging from age 18 to 90. The relationships between SA, TA, and graph metrics were unchanged despite the large variation in age (Figure S1q-t), and our surrogates still captured graph metrics (Figure S7g-i). Since motion was highly correlated with age in this dataset (Figure S3e), we performed all analyses using partial correlation controlling for motion. Several weighted and unweighted graph metrics are correlated with age (Figure S3), with global efficiency, var-FC, and kurt-FC showing significant correlations. Age was positively correlated with SA-λ, negatively correlated with SA-∞, and uncorrelated with mean TA-Δ_1_ (Figure 3a).

**Figure 3.**
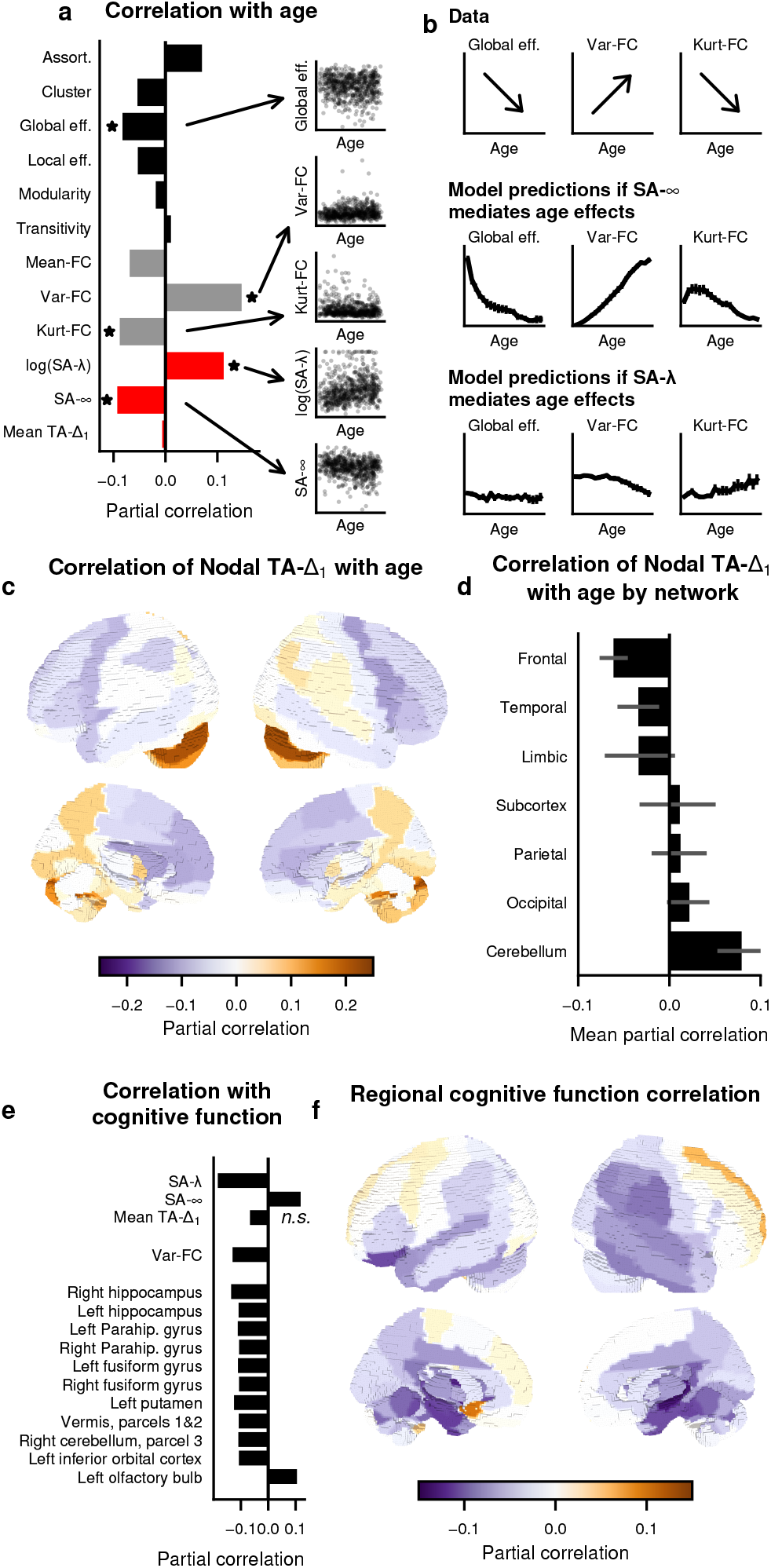
Spatial autocorrelation links functional connectome topology to neurobiology during aging. (a) Partial correlation of graph metrics with age is shown, controlling for motion. Asterisks indicate significance of partial correlation (p<.05). (b) The spatiotemporal model can be perturbed to test whether each SA parameter mediates aging. (top) Subject data predict global efficiency and kurt-FC decrease with age, whereas var-FC increases with age. The SA-∞ and SA-λ parameters were separately perturbed in the spatiotemporal model according to how they change with age. The predicted change of each metric with age is shown if the age-related changes are due to SA-∞ alone (middle) or SA-λ alone (bottom). (c) Partial correlation of TA-Δ_1_ with age is shown for each region, partial on motion. (d) Mean partial correlation of TA-Δ_1_ with age across brain regions, partial on motion. (e) Partial correlation of measures with the ACE-R assessment, partial on movement and age. Top: Partial correlation with SA and mean subject TA. Bottom: all significant partial correlations with mean regional TA-Δ_1_ across subjects. (f) Partial correlation of TA-Δ_1_ with the ACE-R total score, partial on age and motion.

Our spatiotemporal model reveals that the impact of aging on network topology is governed by SA-∞. Since SA-∞ decreases with age, we perturbed the model by decreasing SA-∞^gen^ to obtain model predictions for SA-∞-mediated aging. We found that the predictions match all significant correlations between age and graph metrics from the data (Figure 3b). By contrast, since SA-λ increases with age, we also perturbed the model by increasing SA-λ^gen^ to obtain predictions for SA-λ-mediated aging. However, these predictions do not match the data (Figure 3b). This suggests that the impact of aging on network structure is mediated by changes in SA-∞, i.e. the amount by which nearby connections are favored over distant connections.

Since the mean TA did not change with age, we asked whether this was also true on a regional level. We computed the partial correlation of age and regional TA-Δ_1_ across the brain for each brain region (Figure 3c). Regions in the frontal subnetwork showed significant decreases in TA-Δ_1_ with age (29% of regions significant, p<.05), and cerebellar regions showed increases with age (44% of regions significant, p<.05) (Figure 3d).

### Subclinical markers of dementia

While SA and TA relate closely to network topology, are they useful in explaining clinical symptoms beyond network topology? We explored the relationship of SA and TA with early symptoms of dementia, asking whether SA or TA predict cognitive decline beyond the effect of healthy aging. To assess cognitive function, we used the Addenbrookes Cognitive Examination Revisited (ACE-R), a battery of cognitive tests for dementia screening in sub-clinical populations. We computed the partial correlation across subjects of ACE-R score with SA-λ, SA-∞, and TA-Δ_1_, after accounting for age and movement. We found a significant negative partial correlation with SA-λ (*r* = −0.184, *p* < 10^−5^), associating wider SA with reduced cognitive function (Figure 3e). There was also a weaker correlation with SA-∞ (*r* = 0.118, *p* = 0.002), but no significant relationship with mean TA-Δ_1_ (*r* = −0.067, *p* = 0.09) (Figure 3e). Of all the graph metrics, only the weighted graph metric var-FC (*r* = −0.13, *p* = 0.001) showed a significant relationship (others, *p* > 0.1). This SA-λ-driven effect contrasts with healthy aging, which was primarily driven by SA-∞.

Given the heterogeneity in regional TA, we also examined whether TA-Δ_1_ in specific regions was more predictive than mean TA-Δ_1_. For each brain region, we computed the partial correlation across subjects of ACE-R score with regional TA-Δ_1_, after accounting for age and movement. Across the entire brain, we found eleven brain regions which showed a significant (*p* < 0.01) partial correlation (Figure 3e). These regions were focused around inferior temporal cortex, including bilateral hippocampus, bilateral parahippocampal gyrus, and bilateral fusiform gyrus among the eleven brain regions which showed significant correlation (*p* < .01) (Figure 3e,f). These findings hint at a relationship of clinical symptoms with SA and TA ^16^.

#### Pharmacological manipulation

Lastly, we tested whether SA and TA causally reflect neurobiological processes. In principle, it is possible that TA and SA are driven exclusively by noise, morphology, or other reliable artifacts, instead of by differences in brain dynamics. To test this, we performed a pharmacological manipulation of a neural circuit in human subjects using psychedelic drugs. In two pharmacological fMRI experiments, we measured changes in SA and TA caused by the serotonin receptor agonists lysergic acid diethylamide (LSD) ^41^ and psilocybin ^42^. In both experiments, subjects were administered drug or placebo on separate days in a double-blind methodology, and rs-fMRI was performed at early and late timepoints after each administration. Consistent with the datasets reported above, we confirmed the same basic relationships between SA, TA, and graph metrics were preserved under the drugs (Figure S9a-e), despite a small sample size less than 3% that of HCP.

We found that both LSD and psilocybin caused robust overall reductions in TA across cortex. LSD and psilocybin reduced cortical TA at both early and late time-points, in regional- and subject-averaged TA (Wilcoxon rank-sum test *p* < 10^−40^ for all conditions) (Figure 4a, S9f). Subject-level drug effects on TA-Δ_1_ were not due to differences in within-scanner motion (Figure S9h), and drugs caused no significant change in motion for most conditions (Figure S9i). In contrast to the robust effects on TA, no effect of drugs on SA and graph metrics was detectable for this sample size (Figure S9g,j).

**Figure 4.**
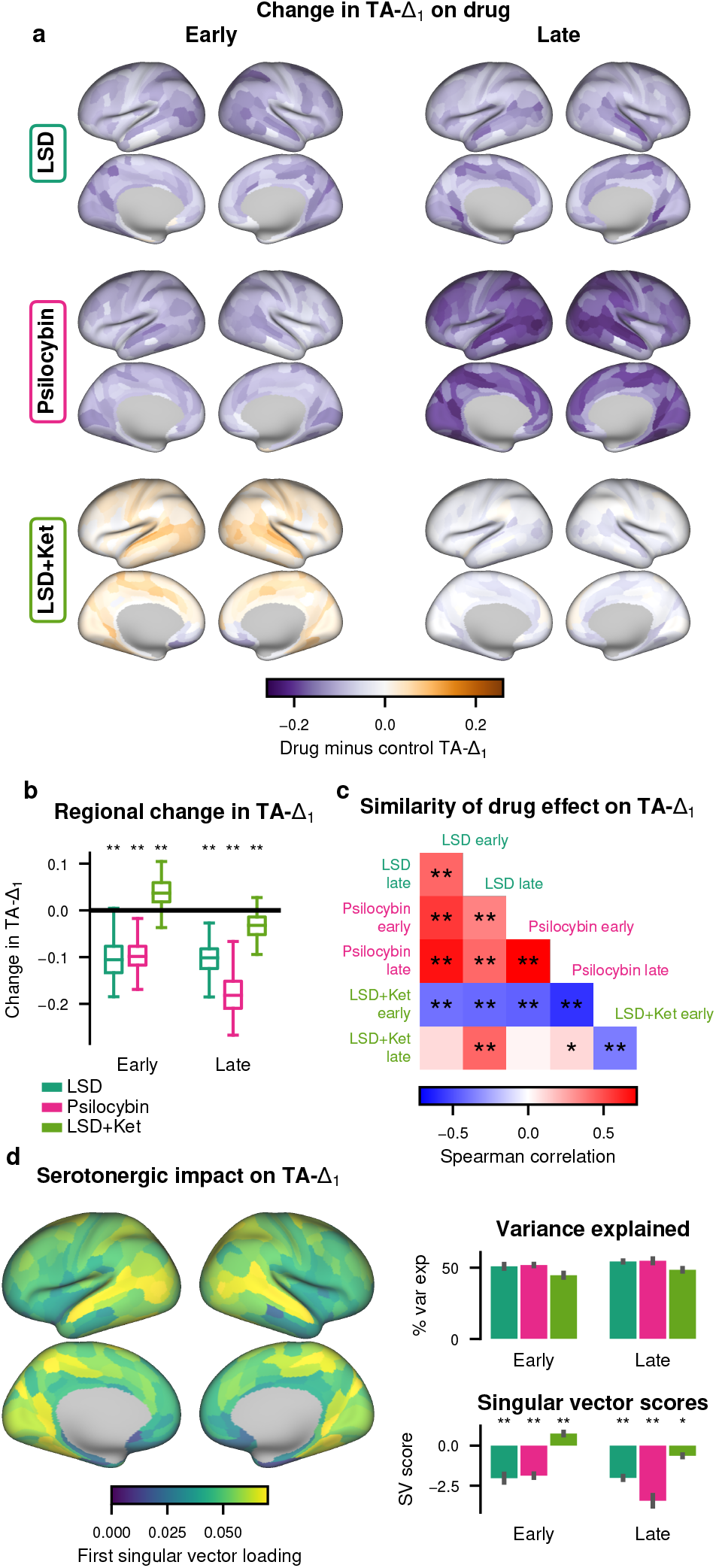
Pharmacological administration of serotonergic psychedelics causes changes in TA. (a) Cortical maps showing the mean difference in TA-Δ_1_ between drug and placebo at early (left) and late (right) timepoints. Purple indicates lower TA-Δ_1_ in the drug than placebo conditions. (b) Change in the regional TA-Δ_1_ for each region, averaged across subjects. (c) Similarity, measured by Spearman correlation, between “drug minus placebo” cortical maps for each drug and timepoint. * indicates p<.05, ** p<.01, two-tailed permutation test correcting for spatial autocorrelation ^8^. (d) The first right singular vector (SV) across all drug minus control contrasts for all subjects. Right top: variance explained by the first SV averaged across each drug condition. Right bottom: the SV score averaged across each condition. Error bars indicate standard error of the mean. * indicates p<.05, ** p<.01, two-sided Wilcoxon signed-rank test.

Psychedelic effects of LSD and psilocybin are pre-dominantly attributed to agonism of serotonin receptors, in particular the 5-HT_2A_ receptor ^41,42^. A common serotonergic mechanism for LSD and psilocybin should produce similar cortex-wide topographies of the change in TA-Δ_1_. Indeed, we found significant positive correlation among the cortical topographies for both drugs and both timepoints (Figure 4c).

Since both of these 5-HT_2A_ agonists produced a specific topographic pattern of reductions in TA-Δ_1_, we predicted that a 5-HT_2A_ antagonist would produce the same pattern but as an increase in TA-Δ_1_ instead of a decrease. In the LSD study, on a third visit, subjects were pre-administered ketanserin, a selective 5-HT_2A_ antagonist ^43^, before receiving LSD (LSD+Ket). We found that at the late timepoint, ketanserin strongly attenuated the effect of LSD on TA-Δ_1_ (Figure 4a). At the early timepoint, there was a cortex-wide increase in TA-Δ_1_ (Figure 4a). Furthermore, regions with the strongest decrease under LSD showed the strongest increase with pre-administration of ketanserin (Figure 4b). These time-dependent changes in TA are consistent with observed pharmacokinetics of ketanserin, which exhibits relatively fast decrease in plasma levels after initial administration ^44^.

We used these experiments to construct a cortical map of overall TA-Δ_1_ modulation by serotonergic drugs. To incorporate information from all experimental conditions, we used singular value decomposition (SVD) to compute the first singular vector across all drug vs control contrasts from all participants. We found a single map (Figure 4d, left) explained approximately 50% of the variance of individual subjects for each of the three experiments at both timepoints (Figure 4d, top right). This map weighted negatively on the LSD and psilocybin conditions and on the late LSD+Ket condition, but positively on the early LSD+Ket condition (Figure 4d, bottom right), consistent with their correlational structure (Figure 4c). These findings demonstrate that TA is sensitive to causal pharmacological perturbation by serotonergic drugs, showing that TA reflects meaningful differences in neurobiology.

## Discussion

Here, we have shown that spatial and temporal autocorrelation—as parameterized by SA-λ, SA-∞, and TA-Δ_1_—are highly reliable properties of rs-fMRI timeseries that correlate with, and are predictive of, network topology. A spatially- and temporally-autocorrelated surrogate timeseries model demonstrated that SA and TA are sufficient to reproduce graph metrics. The surrogate timeseries model was also used to track age-related changes in graph metrics. SA and TA correlate with subclinical symptoms of dementia, and pharmacological manipulations with serotonergic drugs modulate TA in a distinct and reliable pattern. The high reliability, interpretability, effect size, and sensitivity to neurobiology makes spatial and temporal autocorrelation serious candidates for fMRI-based biomarkers ^16,36^.

What factors drive SA and TA? We showed SA and TA can be influenced by biological factors, including aging; pharmacological factors, such as serotonergic drugs; and methodological factors, such as motion, parcel size, and noise. These findings are consistent with previous work in brain signal variability during aging ^45^, as well as in non-serotonin neuromodulators such as norepinephrine ^7,46^, dopamine ^47^, and acetylcholine ^48^. They are also consistent with methodological studies, showing the statistical considerations of SA and TA ^28,49,50^ and the ability of null models to form complex networks ^32,51^. Furthermore, SA and TA are influenced by local circuit connectivity and properties such as intrinsic timescale ^9,16,17^. The influence of confounding physiological processes on SA and TA must be better understood ^52–54^, including the link between TA and signal variability ^55^, especially related to aging ^45^. The hemodynamic response function imposes not only temporal but also spatial filtering to the BOLD signal ^56,57^, which could contribute to changes in SA and TA due to age ^58^ or serotonergic drugs ^59^. Since anatomical structure serves as a scaffold for SA and TA ^10,11,16,33^, individual differences in brain morphology, as measured by structural or diffusion MRI, might explain our excellent test-retest reliability and fingerprinting performance, offering insights into the relationship between brain structure and function ^8,16,30,52,60^.

In general, SA and TA constrain the dimensionality of the neural signal, and these dimensionality constraints may explain their link to network topology. Previous work has shown lower-dimensional subspaces capture network topology ^22–24^, and we showed SA and TA align with these dimensions through the economic clustering model. These dimensions may map onto other aspects of network topology, e.g., a high degree backbone and a lattice-like background ^40^. The eigensurrogate model tested the broader question of whether another FC matrix with equivalent linear dimensionality could reproduce graph metrics. While it did not perform as well as our timeseries-based model on most datasets, it performed surprisingly well and even outperformed our model on the Yale-TRT dataset, emphasizing the role of constrained dimensionality in shaping network topology.

Our present study used a parcellated analysis, which limits the precision with which SA can be measured. Our methods for computing both SA and TA scale well for voxel- or vertex-level analyses, but fitting the spatiotemporal model is computationally intractable for parcellations with many nodes. SA-∞ is conceptually similar to mean-FC, which has known links to aging ^52,61^. It is also similar to global signal, but it remains reliable even after global signal regression, and thus may represent spatial inhomogeneities in the global signal. Nevertheless, even this parcellated analysis is sensitive enough to reveal features inaccessible to graph theoretical analysis. For example, in our pharmacological experiments, our sample size of less than 25 participants was too small to show significance in graph-theoretical measures, but changes in TA revealed highly significant differences in functional organization. Our approach only coarsely matches SA by using Euclidean distance to fit subject-level parameters, limiting its ability to capture nodal properties. Future work may adapt our measure of SA to include spatial inhomogeneities in SA, or to allow alternative measures of distance such as geodesic distance in a single hemisphere.

While we focus here on how SA and TA impact network topology using graph theory, all analyses of resting-state FC may be influenced by SA and TA ^8,16,30^, including task-related changes ^12,13^. The impact of SA and TA on rs-fMRI analysis can be mitigated in a number of ways. Main effects can be analyzed for a correlation with SA and TA to show if they are related. While the effects of SA and TA are nonlinear, adjusting for these factors using techniques such as partial correlation can also reduce their effect. Reporting confounds of experimental methodology and preprocessing on SA and TA allows their influence to be summarized. Likewise, spatially-^8,25,26^ and temporally-informed ^32^ null models allow for robust statistical analyses despite the presence of SA and TA. However, our results overall suggest that perhaps SA and TA should not be treated as confounds, but rather, as essential informative properties of the human connectome.

## Acknowledgments

We thank Josh Burt for help with plotting; Amber Howell for assistance with HCP data; and Taku Ito and Jean C.C. Vila for helpful discussions. Funding was provided by the European Molecular Biology Organization grant ALTF 712-2021, the Winston Churchill Foundation of the United States, and the Gruber Foundation (MS); NIMH grant K00MH122372 (SN); Swiss National Science Foundation (P2ZHP1\161626) (KHP); Agence Nationale de la Recherche (ANR-20-NEUC-003-01) and MIAI@Grenoble Alpes (ANR-19-P3IA-0003) (SA); Usona Institute (2015-2056), Heffter Research Institute (1-190420), Swiss Neuromatrix Foundation (2015-103 and 2016-0111), and Swiss National Science Foundation under the framework of Neuron Co-fund (01EW1908) (FXV); NIHR (Senior Investigator award) and the NIHR Cambridge Biomedical Research Centre (ETB); SFARI (Pilot Award) (AA and JDM); and the National Institute of Mental Health (NIMH) (R01MH112746) (JDM). Data collection and sharing for this project was provided in part by the Cambridge Centre for Ageing and Neuroscience (CamCAN). Cam-CAN funding was provided by the UK Biotechnology and Biological Sciences Research Council (grant number BB/H008217/1), together with support from the UK Medical Research Council and University of Cambridge, UK.

## Author contributions

MS and ETB conceived the research. MS designed the experiments. MS and AH performed the experiments. MS and JDM analyzed and interpreted results. KHP, JLJ, FM, SN, DS, RTC, JHK, FXV, and AA contributed data, methodology, and resources. MS, LT, and SA performed the mathematical analysis. DL, ETB, and JDM provided supervision and funding. MS, AH, and LT wrote the first draft of the manuscript. All authors edited, revised, and approved the manuscript.

## Methods

### Datasets

We analyzed the following datasets, which comprise a diversity of preprocessing steps and experimental methodologies, including different parcellations, sampling rates, spatial and temporal smoothings, covariate regressions, and noise removal strategies.

#### Human Connectome Project (HCP)

A total of 883 subjects aged 22-37 from the Human Connectome Project 1200 subject data release underwent four resting-state scanning sessions spread across two days. Resting-state scans lasted for 14.4 minutes with a TR of 0.72 s (sampling rate of 1.39 hz). Data were preprocessed with the Human Connectome Project minimal preprocessing pipeline. This includes distortion correction using the field map, realignment for head motion, registration to T1 images. Data were further denoised using ICA-FIX, a technique based on independent component analysis to remove structured noise. A high-pass filtered was applied at 0.01 hz. Data were parcellated into 360 regions (180 per hemisphere) ^18^ with multimodal surface matching based on MSMAll ^63^, and with 2 mm FWHM surface spatial smoothing constrained to the parcel. The first 100 timepoints were removed to ensure steady state.

#### Human Connectome Project with Global Signal Regression (HCP-GSR)

Subjects and scan parameters are identical to the HCP dataset, but global signal regression was added as a preprocessing step before parcellation. Additionally, we did not truncate the first 100 TRs, for a total of 1200 timepoints. 33 additional subjects were excluded due to nonconvergence in the global signal regression pipeline for a total of 850 subjects.

#### Yale Test-Retest

A total of 12 subjects were scanned on four different days, and six 6-minute sessions were performed each day. Each subject was scanned on two scanners, two days on each scanner, on days spaced approximately one week apart. Thus, half of the scanning sessions for each subject were performed on a different scanner. Scanning sessions lasted 6 minutes each with a TR of 1.0 (sampling rate 1.0 hz). Motion correction was applied, and images were spatially smoothed to achieve uniform spatial smoothness of a 2.5mm Gaussian kernel ^64,65^. Images were coregistered to a common subject-specific space across days, and subsequently into MNI space, and parcellated using the Shen parcellation ^66^.

High pass filtering was performed by regressing out linear, quadratic, and cubic trends, and low pass filtering was performed with a Gaussian kernel with cutoff frequency of 0.19 hz. Mean white matter, mean cerebrospinal fluid, mean global signal, and a 24-parameter motion model were also regressed out of the data. No subjects, sessions, or regions were excluded.

#### Cambridge Centre for Ageing and Neuroscience (Cam– CAN)

A total of 652 subjects were scanned using a TR of 1.97 s (sampling rate 0.508 hz). We utilized the standard preprocessing pipeline and parcellation provided by the Cam-CAN project ^20,68^. In summary, scans underwent motion correction and slice time correction before coregistration to T1 images and normalization to MNI space with DARTEL. We utilized the default AAL parcellation provided by the Cam-CAN project ^69^. We also applied a second-order Butterworth low-pass filter at half the Nyquist frequency (0.127 Hz) to account for high-frequency motion artifacts. We excluded six subjects and one cerebellar region due to missing data, for a grand total of 646 subjects with 115 regions in the parcellation.

#### LSD

The study utilized a double-blind randomized design ^41^. On each of the three days, participants received one of the following treatments: (a) placebo (179 mg Mannitol and 1 mg Aerosil, orally) pretreatment followed by placebo (179 mg Mannitol and 1 mg Aerosil, orally) treatment; (b) placebo (179 mg Mannitol and 1 mg Aerosil, orally) pretreatment followed by LSD (100 μg, orally); and Ketanserin (40 mg, orally) pretreatment followed by LSD (100 μg, orally) treatment. Pretreatment was given 60 minutes before the treatment. The “early” resting state scan occurred 75 min after treatment, and the “late” scan 300 min after treatment. Data were acquired with TR=2500 ms (sampling rate 0.4 hz). 25 subjects were enrolled in the study. One subject was excluded for technical faults in the data for a total of 24 subjects.

We utilized the same preprocessing pipeline described in ^41^, which is summarized below. First, the data were subject to the HCP minimal preprocessing pipeline. This involved correction for field inhomogeneities, phase encoding directions, and magnetic susceptibility artifacts, as well as motion correction and registration to structure images with non-brain tissue masking. Data were high pass filtered (>.008 hz). Several nuisance variables were regressed out: average signal in the ventricles, and in deep white matter, motion parameters, and the mean timeseries across gray matter (i.e. the global signal), as well as the first derivative of each of these. Data were motion scrubbed, identifying outlier frames as either frames with a summed framewise displacement over 0.5 mm, or a RMS of differences in intensity of subsequent frames over 1.6 times the median. All outlier frames, the one frame preceding them, and two frames following them, were excluded from analysis. Lastly, data for cortex were parcellated into 360 regions according to the Glasser parcellation ^18^. Measurements of TA-Δ_1_ were performed only on consecutive segments involving no dropped frames, such that TA-Δ_1_ was a weighted average of the TA-Δ_1_ of each consecutive segment based on the degrees of freedom of that segment. The long distance correlation parameter *d* was not computed on scrubbed data.

#### Psilocybin

The study utilized a double-blind randomized design ^42^. Participants were scanned two separate days. On each day, participants received either placebo (179 mg Mannitol and 1 mg Aerosil) or psilocybin (0.2 mg/kg), administered orally. rs-fMRI was performed at three timepoints following psilocybin administration: “immediate”, performed 20 minutes after administration, “early” after 40 minutes, and “late” after 70 minutes. Data were acquired with TR=2430 ms. 24 participants were enrolled in the study. One participant was excluded due to a missing scan, for a total of 23 participants. This sample size was determined based on the prior LSD study ^41^.

Data preprocessing was identical to that of the LSD dataset described above.

### Estimating spatial autocorrelation

We decomposed spatial autocorrelation into two components: the rate at which correlations decrease exponentially with distance (SA-λ), and the spatially-invariant level of correlation to which it decays (SA-∞). This can be described quantitatively as

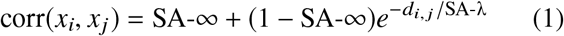

where *x*_*i*_ is the timeseries for region *i, d*_*i, j*_ is the Euclidean distance between regions *i* and *j*, and corr is the Pearson correlation. The quantities SA-λ and SA-∞ are constants and do not depend on *i* or *j*.

We estimated the parameters SA-λ and SA-∞ for each subject as follows. For each pair of brain regions, we computed both their physical Euclidean distance (from the region’s centroid) as well as their Pearson correlation coefficient. We used Euclidean distance rather than geodesic distance because it is defined across cortical hemispheres and within subcortex. We binned each pair of brain regions according to their Euclidean distance, using 1 mm bins for the HCP and HCP-GSR datasets, and 5 mm bins for the TRT and Cam-CAN dataset due to the fewer number of parcels, and computed the mean Pearson correlation of pairs in each bin to generate correlation vs distance curve (Figure 1a). We then found the least square fit of Equation 1 to this curve, optimizing with gradient descent, bounding SA-λ between 0 and 100 and SA-∞ between -1 and 1. While it is possible to fit Equation 1 directly to the distance and correlation of each pair without binning, our approach puts more weight on nearby and distant correlations, which are less represented in the data but most critical for determining SA-λ and SA-∞.

To account for heteroskedasticity in SA-λ for the Cam-CAN dataset, we analyzed the logarithm of this parameter for this dataset.

### Estimating temporal autocorrelation

Our primary measure of temporal autocorrelation, TA-Δ_1_, is a non-parametric measurement computed by taking the Pearson correlation of neighboring timepoints in the parcellated timeseries, i.e., for a timeseries *x* [*t*], we have corr (*x* [*t*], *x* [*t* + 1]). This measure is computationally efficient, and can be implemented with only a few lines of code. However, TA-Δ_1_ is not comparable across datasets due to differing TR.

### FC and graph construction

We defined the FC matrix ρ_*i, j*_ as the matrix of Pearson correlation coefficients ρ_*i, j*_ = corr (*x*_*i*_, *x*_*j*_) between each pairwise combination of regional timeseries *x*_*i*_ and *x*_*j*_.

We constructed unweighted, undirected graphs from model or data FC matrices using standard techniques ^4,5^. We first constructed a spanning tree backbone to ensure connectedness by transforming each element of the FC matrix ρ_*i, j*_ by 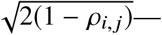an operation which turns the correlation similarity measure into a distance ^74^—and then applying Kruskal’s algorithm to find the minimum spanning tree ^129^. We iteratively added the strongest edges in the FC matrix to the spanning tree until the graph contained 10% of all possible edges (proportional thresholding). This produced an unweighted, undirected graph with a fixed number of edges.

### Quantifying reliability

We quantify univariate reliability using the intraclass correlation coefficient (ICC). ICC measures the reliability of a particular scalar measure across subjects ^38,76^. Let *N* be the total number of subjects, *M* be the number of sessions per subject, and 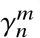 be some scalar measure of interest for session *m* of subject *n*. Similar to an ANOVA, ICC decomposes the variance across subjects into variance from a common source, 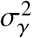, and noise, *σ*_*E*_, by assuming that γ can be decomposed into a subject-specific term γ_*n*_ and a noise term E_*n,m*_, i.e., 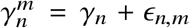. Then, the ICC is defined as the fraction of variance explained by the subject-specific term, i.e.,

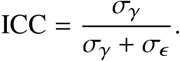

In theory, there are generalizations of the ICC which can accommodate additional sources of variance. Here, we use the simplest form, ICC(1,1) ^38^.

We quantify multivariate reliability using fingerprinting, similar to that performed in Ref. ^19,37,80^. Let 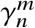 be a vector describing some measure (e.g. TA-Δ_1_) from a single session *m*∈ {1 … *M*} for subject *n* ∈ {1 … *N*}. For each session *m* of each subject *n*, we compute

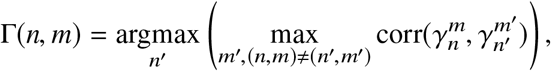

where corr is Pearson correlation. The fingerprinting performance is given by

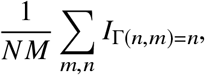

where *I* is the indicator function. Under this measure, chance performance is (*M* – 1) (*N*− *M* −1). For regional measures, the length of vector γ^*m*^ was equal to the number of regions in the parcellation *R*. For FC, the length of 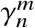 was *R* (*R* −1) /2, i.e. the number of distinct elements in the FC matrix.

In the HCP and HCP-GSR datasets, subjects were scanned four times over two days. For computing ICC, we used *M* = 4. For fingerprinting, in order to obtain error estimates and increase the difficulty of fingerprinting, we performed fingerprinting on each possible pair of sessions 1-4 (*M* = 2, performed six times, instead of *M* = 4 performed once). This means each session had only one correct match of the 1765 sessions in the pool. In the TRT dataset, subjects were scanned in six sessions across four different days for a total of 24 sessions per subject. Likewise, we used *M* = 24 to compute ICC. By contrast, we performed fingerprinting for each session independently (i.e. *M* = 4 performed six times instead of *M* = 24 performed once), so subjects had 3 correct matching sessions out of 47 in the pool. Without such increases in difficulty on these datasets, fingerprinting performance was near 100%. We could not compute ICC or perform fingerprinting on the Cam-CAN dataset, since only one rs-fMRI session was acquired per subject.

### Weighted graph metrics

We consider three weighted graph metrics: the mean (mean-FC), variance (var-FC), and kurtosis (kurt-FC) of the FC matrix. Each is calculated by finding the corresponding statistic (mean, variance, or kurtosis) of the upper triangular portion of the FC matrix, excluding the diagonal. In other words, for the FC matrix *M*, this is the mean, variance, and kurtosis of the set ℳ={*m*_*i, j*_ : *i* < *j*}.

Note that the thresholding procedure destroys all explicit information about mean-FC, var-FC, and kurt-FC. To understand why, recall that the thresholding procedure keeps a fixed fraction of edges. Thus, if we threshold an FC matrix, the number of edges only depends on the fixed fraction of edges we keep. For an FC matrix of size *N*, if we keep *q* fraction of edges, then the moments are identical to those of the binomial distribution: mean-FC of the thresholded matrix is *q*, the variance is *q* (1−*q*), and the kurtosis is 1/*pq* − 6. Therefore, no explicit information remains about mean-FC, var-FC, or kurt-FC after thresholding.

Because TA can be measured for individual nodes, we also consider how it impacts local topology. To do this, we also define the corresponding nodal graph metrics for each which operates on rows of the FC matrix instead of the upper diagonal: nodal mean-FC, the mean of the row; nodal var-FC, the variance of the row; and nodal kurt-FC, the kurtosis of the row. We exclude self-connectivity (i.e. the diagonal of ones in the FC matrix) from these calculations. Thresholding does not destroy nodal mean-FC, nodal var-FC, or nodal kurt-FC.

### Unweighted graph metrics

We consider six popular graph metrics to quantify the topology of the connectome ^6^:

#### Assortativity

Assortativity is a preference of high-degree nodes to connect to each other. Mathematically, this is the Pearson correlation between the degree of nodes connected by edges ^82^.

#### Clustering coefficient

The clustering coefficient for a single node is the average relative number of triangles around a node. For adjacency matrix *A*, the nodal clustering coefficient for node *i* is

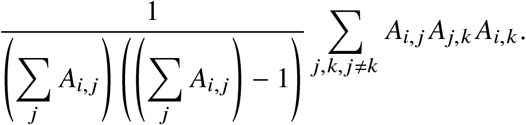

The clustering coefficient for the network is the average of the nodal clustering coefficients.

#### Global efficiency

The global efficiency is related to the average topological distance between nodes. Mathematically, it is the mean of the inverse of shortest path lengths between each pair of nodes, i.e.

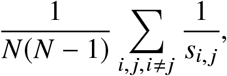

where *s*_*i, j*_ indicates the shortest path between nodes *i* and *j* and *N* is the number of nodes ^83^.

#### Local efficiency

Local efficiency, similar to clustering, is the mean global efficiency on the subgraph of each node’s nearest neighbors ^83^.

#### Modularity

Modularity quantifies the ability to break a network into “communities” such that the number of edges within the community is maximized and outside the community is minimized. Let *C*_*i*_ be some community assignment of node *i*, and *δ* be the Kronecker delta function. We can compute the quality of the community assignment *C*_*i*_ with the equation

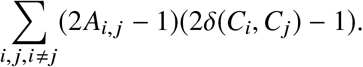

The first term in the sum is 1 if the neurons are connected and -1 if they are not, and the second term is 1 if they are in the same community and -1 if they are not. Thus, this can be maximized if connected nodes fall into the same community and unconnected nodes do not. The modularity is defined as the maximum value of this function across all potential community assignments {*C*_*i*_}_*i*_, rescaled to fall between -0.5 and 1^84^

#### Transitivity

The transitivity is the total number of number of 3-way reciprocal connections compared to the total possible number of such connections, i.e.,

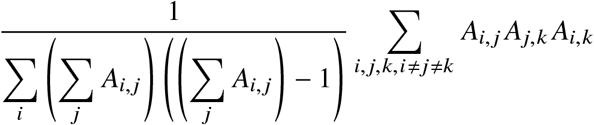

Note that this is distinct from local efficiency because it considers the network as a whole rather than considering each node *i* individually and then averaging.

In addition to considering these graph metrics, we also consider two nodal graph metrics:

#### Degree

The degree is a nodal metric which measures the total number of edges connected to a node, i.e.,

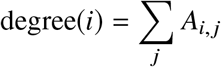

#### Centrality

Betweenness centrality is a nodal metric which measures the faction of shortest paths between all pairs of nodes in a network which pass through the given node. It does not have a closed form equation and must be computed using an algorithm ^85^.

### Graph metrics linear model

To quantify the impact of SA and TA jointly, we utilized a linear model. We fit the “SA-λ + SA-∞” model

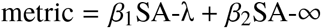

or the “All” model

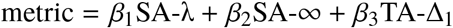

on 50% of the data, randomly chosen from all sessions. Shown in Figure 1e and Figure S1 is the Spearman correlation of the 50% held-out data with the predictions of this linear model. We used Spearman correlation instead of *R*^2^ for fair comparison SA-λ, SA-, and TA-Δ_1_ individually. Because the fit uses held-out data, the correlation with the linear model does not necessarily need to be higher than the correlation with any of the individual factors in the model.

### Spatiotemporal model

The spatiotemporal model generates surrogate timeseries which can be analyzed like rs-fMRI timeseries. It takes two parameters— the noiseless SA-λ (SA-λ^gen^), and the noiseless SA-∞(SA-∞^gen^)—and also uses two pieces of information from the data—the TA-Δ_1_ *ϕ*_*i*_ from each region *i*, and the Euclidean distance *d*_*i, j*_ between the centroids of each pair of regions *i* and *j*.

The model operates in two basic steps. First, we generate random timeseries which are both spatially and temporally autocorrelated. For this first step, all timeseries have uniformly high TA, and SA is determined by the parameters SA-λ and SA-∞. Using the derivation outlined in Supplement 2.1, we generate *N* high-pass filtered (cutoff frequency 0.01 hz, 4th order Butterworth filter) Brownian noise timeseries (frequency spectrum 1 / *f* ^2^) of length *T* such that, for timeseries *x*_*i*_ [*t*] and *x*_*j*_ [*t*], we have

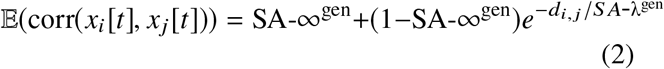

where corr is the Pearson correlation, similar to Equation 1. The matrix consisting of all such expected correlations between regions *i* and *j* forms the matrix *C*_*i, j*_ required by the algorithm described in Supplement 2.1, such that

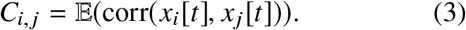

For non-negative SA-∞^gen^, *C*_*i, j*_ will be positive semidefinite. Second, we reduce the temporal autocorrelation of each timeseries to match that of the original data. Brownian motion timeseries have high TA and adding white noise to a timeseries reduces TA, so we add white noise to each timeseries *x* [*i*] until the TA-Δ_1_ of the timeseries is equal to the empirical TA-Δ_1_ *ϕ*_*i*_. Thus, we have to choose a distinct amount of noise to add to each timeseries such that corr *x*_*i*_ [*t*], *x*_*i*_ [*t* + 1] = *ϕ*_*i*_. We update *x*_*i*_ [*t*] *← x*_*i*_ [*t*] + *N* (0, *h* (*ϕ*_*i*_)) for some function *h* (*ϕ*_*i*_) given by Equation 4 using the analytical derivation outlined in Supplement 2.3. The function *h*(*ϕ*_*i*_) is defined for *ϕ*_*i*_ *>* 0, and the variance of a random variable must be non-negative, so *ϕ*_*i*_ is truncated such that *ϕ*_*i*_ ≥ 0.0001 and *h* (*ϕ*_*i*_) ≥ 0. Note that SA-λ^gen^ and SA-∞^gen^ are parameters of the underlying process, and differ from the SA-λ and SA-∞ of the generated timeseries due to the addition of noise. The resulting timeseries exhibit a spatial embedding given by the parcel centroid distances *d*_*i, j*_ and parameters SA-λ^gen^ and SA-∞^gen^, and have regional TA-Δ_1_ equal to the original timeseries.

### Model variants

We test several variants of the model to determine the importance of three key components of the model: added uncorrelated noise, TA, and SA.

#### TA only

This model modifies the spatiotemporal model to remove the spatial embedding. Specifically, it fixes the parameters SA-λ^gen^ and SA-∞^gen^ to 0, such that corr(*x*_*i*_ [*t*], *x*_*j*_ [*t*] = *I*_*i*=*j*_ where *I* is the indicator function. It takes no parameters.

#### SA only

We generate random multivariate Gaussian noise with mean zero and covariance matrix

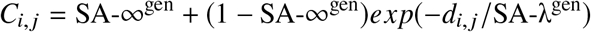

similar to Equation 1. It takes two parameters, SA-λ^gen^ and SA-∞^gen^. Unlike the spatiotemporal model, E(SA-λ) = SA-λ^gen^ and E(SA-∞) = SA-∞^gen^.

#### Noiseless

In our spatiotemporal model, we make sure timeseries have the desired TA-Δ_1_ by generating timeseries with uniformly high TA-Δ_1_, and then adding different magnitudes of white noise to each timeseries to match TA-Δ_1_ to the original timeseries. In this model, we do not add white noise to the timeseries. To match TA-Δ_1_ to the original timeseries, we generate timeseries directly with matched TA-Δ_1_. This is possible because the derivation presented in Supplement 2.1 does not require each timeseries’ power spectrum to be identical. Thus, we achieve this diversity in TA-Δ_1_ by assuming each timeseries has a filtered pink noise (1 / *f* ^α^) temporal dynamics. Then, for each timeseries *i*, we find an exponent *α*_*i*_ such that the high pass filtered 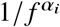 spectrum has expected TA-Δ_1_ *ϕ*_*i*_.

Thu success of this procedure requires us to choose *α*_*i*_ such that the high pass filtered 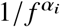 spectrum has TA-Δ_1_ equal to *ϕ*_*i*_. The mapping *ϕ*_*i*_ → *α*_*i*_ can be determined by numerically inverting the *α*_*i*_ → *ϕ*_*i*_ mapping implied by Supplement 2.2. High pass filtering is performed at the level of the power spectrum by multiplying the square of the amplitude response of the filter by the power spectrum. Additionally, in this model, it is possible to determine the SA-λ^gen^ and SA-∞^gen^ parameters directly from the data, without the need to fit parameters, by using the procedure is outlined in Supplement 2.1. As in the “SA only” model, 𝔼 (SA-λ) = SA-λ^gen^ and 𝔼 (SA-∞) = SA-∞^gen^.

Thus, in summary, this model makes the following modifications to the spatiotemporal model: (1) rather than simulating random walks (1 / *f* ^α^ where *α* = 2), the spectral exponents *α*_*i*_ for each region *i* are chosen such that the resulting TA-Δ_1_ is equal to each region’s TA-Δ_1_ value; and (2) no noise is added to the powerlaw timeseries.

#### Homogeneous

*TA-*Δ_1_ To allow SA and TA to be independently manipulated, we developed a homogeneous variant of the spatiotemporal model which treats TA as a parameter. Rather than use TA-Δ_1_ values computed from the original timeseries, this model uses a single fixed value of TA-Δ_1_, 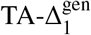, for all regions. For simplicity, we fixed SA-∞^gen^ to be the mean SA-∞ across all networks. Thus, the model takes two parameters: 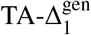 and SA-λ^gen^.

### Eigensurrogate model

Due to our use of eigenvalues for fitting, we developed the eigensurrogate model to test whether eigenvalues alone are capable of reproducing a phenomenon. This tests the null hypothesis that an effect can be explained by its linear dimensionality. Unlike most of the other models we considered, this produces surrogate FC matrices instead of surrogate timeseries.

We first performed an eigendecomposition of the correlation matrix (FC matrix), and then applied the procedure of Ref. ^86^. Briefly, we sampled a random set of eigenvectors, and then applied a series of rotations to set ones on the diagonal. We used the method as implemented by Scipy in the numerical Python stack.

Since this model creates surrogate FC matrices, we produced timeseries by sampling the maximum entropy timeseries which would produce such a correlation matrix. To do this, we numerically computed the matrix square root and multiplied it by a *N* × *T* matrix of standard normal iid random variables. This is equivalent to sampling each timepoint independently from a multivariate Gaussian distribution. In principle it is possible to create temporally-autocorrelated timeseries from this model (multiplying by temporally-autocorrelated timeseries instead of iid standard normal random variables), but by definition, any method of generating timeseries from the eigensurrogte method will produce an identical FC matrix.

### Null models

We also test several popular null models.

#### Phase randomization

The power spectrum amplitude of individual timeseries is preserved, but the phases are randomized (Figure S4). This procedure is described in detail in ^87^. A Fourier transform was performed on each region’s timeseries. Each element of the complex-valued Fourier transform was randomly rotated on the unit circle, and the inverse transform was performed on the phase-randomized spectra.

Note that we used distinct random phases for each timeseries, contrary to many neuroimaging studies which use the same phase for a given frequency across all timeseries. This is because using the same phase for each timeseries preserves all cross-correlations between timeseries. In practice, this means that the surrogate FC matrix is identical to the original FC matrix, and hence, it produces a identical graph.

#### Zalesky matching

This model matches the mean and the variance of the correlation matrix, and is described in full in ^32^. Briefly, it matches the first two moments by iteratively computing correlation matrices from timeseries of different durations with different ground-truth correlations. The process continues until the timeseries duration is found which maximally reproduces the mean and variance of the correlation matrix.

#### Edge reshuffle

We preserve the degree of each node while scrambling the edges (Figure S4). This is accomplished by an iterative algorithm described in detail in ^89^. Two edges from a graph were selected at random, and the connections were swapped. This swap was iterated *k* times, where *k* was chosen here to be 5 times the number of edges in the network. The result is that each node has the same number of connections, but those connections are randomized.

### Models not considered

Despite our parameterization based on the TA-Δ_1_ we do not report on timeseries generated using autoregressive (AR) or vector autoregressive (VAR) models. These models have two limitations within this context. First, they do not reproduce the observed long-memory processes observed within rs-fMRI timeseries. Second, when fitting data using these models, the TA-Δ_1_ parameter fits to values very close to 1, resulting in parameter degeneracy.

Likewise, we did not directly fit the economical clustering model to data. Due to the variability in individual instantiations of this model and the lack of smoothness of the parameter space, this model incompatible with our numerical fitting algorithm, thus preventing the use of comparable methodology to perform the fitting. These issues, combined with the long execution time of the model, made such individual level fitting infeasible.

We did not consider models in which only one of SA-λ and SA-∞ is fit and the other is fixed. These two parameters were fit by optimizing a function to the subject’s measured SA. However, we were unable to get consistent and interpretable fits to this function when only one of these two parameters was fit. In other words, fixing one of these parameters precludes reliable estimation of the other.

Lastly, we only considered approaches which operate at the level of the parcellated timeseries. This excluded approaches such as scrambling using a 3D Fourier transform.

### Model fitting procedure

Due to the fact that uncorrelated random noise was added after the spatial embedding in our model, the SA-λ^gen^ and SA-∞^gen^ parameters in the spatiotemporal model were not identical to the observed SA-λ and SA-∞. We derived a mathematical method for directly matching the spatiotemporal model’s SA-λ^gen^ and SA-∞^gen^ parameters to the data without the use of fitting, but technical constraints prevented an implementation of the procedure (Supplement 2.4). Therefore, we could not directly estimate the model based on observed SA-λ and SA-∞.

Instead, the spatiotemporal model’s SA-λ^gen^ and SA-∞^gen^ parameters were fit to individual subject FC matrices to match that matrix’s eigenvalue distribution. The model was fit using differential evolution ^90^, a gradient-free global heuristic search method, to the mean squared error between the sorted eigenvalue distributions of the subject and model FC matrices. Eigenvalues were non-negative due to the positive-semidefiniteness of the correlation matrix. The model was implemented in such a way that they preserved smoothness with respect to the parameters, meaning that for a given random seed, small perturbations of the parameters caused only small changes in the timeseries, and hence in the structure of the graph. Optimization was performed on the mean objective function from two random seeds. All reported statistics and metrics about the models come from a single instantiation of the model using a different random seed than either seed used during fitting. Parameters for the “SA only” model were also fit using the same procedure.

Most alternative models had no parameters (“TA only”, “Phase randomization”, “Zalesky matching”, and “Edge reshuffle”). For the “Noiseless” model, parameters could be fit directly using the formalism in Supplement 2.1. This formalism is not guaranteed to converge for all subjects. When the algorithm was unable to produce valid timeseries, we excluded these subjects from the analysis. Results were qualitatively similar when parameters were fit to the eigenvalue distribution as described above which forced parameters into valid regimes.

### Economical clustering model

Here, our model produces timeseries for each brain region—graphs can then be constructed by processing these timeseries the same way as subjects’ rs-fMRI timeseries. In the graph theory literature, the term “generative model” usually refers to models which construct graphs directly through the iterative addition of nodes or edges ^22,23,119^. The economical clustering (EC) model is one such model which is popular for studying brain networks ^22,23^ (Supplement 4.1). In this model, connections between nodes are determined by one parameter governing the impact of distance and one for the impact of clustered topology. The probability of an edge forming between two brain regions is proportional to the product of the Euclidean distance between the regions raised to some power (the distance parameter), and the fraction of shared neighbors between them raised to some power (the clustering parameter). Full model details are provided in Supplement 4.2 and in Ref. ^22^.

To compare the two models, we simulated the EC model across a spectrum of distance and clustering parameters, and then fit a spatiotemporal model to the simulated networks. Because our spatiotemporal model takes two SA-related parameters and obtains TA on a regional level directly from the data, we compared the EC model to the homogeneous variant of the spatiotemporal model, which includes one SA parameter and one TA parameter. We used the EC model to simulate 10 networks per combination of parameters, fitting the homogeneous spatiotemporal model to each of these 10 networks. Since the EC model produced graphs rather than FC matrices, we could not use our previous approach of fitting by eigenvalues, nor could we derive an analytic approach to parameter estimation. Thus, we fit using the objective function from Ref. ^23^. Full details are provided in Supplement 4.3.

### Spatial correction for brain map similarity

To assess the similarity between brain maps, the presence of spatial autocorrelation can induce a high false positive rate. To correct for this, we perform a permutation test using an SA-preserving surrogate method ^8^. To compare a target brain map to a reference map, we generate 10000 surrogate brain maps which match the spatial autocorrelation of the reference map. Then, we compute the Spearman correlation between the target map to the reference map, as well as between the target map and the surrogate maps. The p-value of the two-tailed test is determined as the fraction of correlations with the surrogate maps which are at least as extreme in absolute value as with the reference map. For cases in which maps cannot be designated as a target or reference map, we perform this procedure twice, once with each map taking the role of the reference map and the other as the target map, and compute the two-tailed p-value as the total number of target-to-surrogate Spearman correlations which as at least as extreme in absolute value as the Spearman correlation between the two maps.

### Singular value decomposition

We computed a cortical map of serotonergic modulation using singular value decomposition (SVD), which bears many similarities to principal component analysis (PCA). For a data matrix M, we can rewrite *M* as

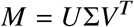

where Σ is a diagonal matrix, and the rows of *U* and *V* form orthogonal bases. The diagonal elements of Σ are called “singular values”, the rows of *V* are called “singular vectors” (each element of which is a “loading”), and the projection of *M* onto *V* (or, equivalently, *U*Σ) are the “scores”. Note also that the product *M*^*T*^ *M* = *V* Σ^2^*V*^*T*^, so Σ^2^ and *V* are the eigenvalues and eigenvectors, respectively, of *M*^*T*^ *M*. If *M* is centered, then *M*^*T*^ *M* is the covariance matrix, and SVD is equivalent to PCA, meaning Σ^2^ gives the variance explained of each component. But in our case, since *M* is not centered, variance explained of the first *k* components is computed as

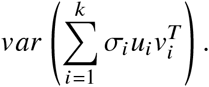

To compute the variance explained by experimental condition, we find the variance explained by each experiment from each subject individually, and then average across the experimental condition.

### Code availability

We released a software package which allows the central analyses in this paper to be peformed quickly and easily, the “spatiotemporal” Python package, which can be installed through pip or downloaded at https://github.com/murraylab/spatiotemporal. All other source code will be made available upon publication. Code was implemented using the standard Python stack ^95,96^ and other libraries ^97–99^. Source code was checked for correctness using software verification techniques ^100^.

## Supplemental information

### Supplement 1: Relationship between TA-Δ_1_ and long memory processes

fMRI timeseries are thought to show characteristics of a long memory process, i.e., time points which are separated by a large lag show a high correlation ^35,101,102^. These processes are often characterized by their autocorrelation function (ACF). For a timeseries *x*, the autocorrelation function is defined as

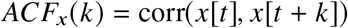

where corr is the Pearson correlation and *k* is the lag at which to evaluate the ACF. Long memory processes have an ACF which decays slowly across lags, meaning that relatively high correlations exist even at large lags. However, in this work, we use TA-Δ_1_ as our measure of temporal autocorrelation, which is identical to the lag-1 term of the ACF, *ACF*_*x*_(1). If fMRI timeseries show long memory properties, and long memory processes are based on many terms of the ACF, this raises the question of whether TA-Δ_1_ is an appropriate measure to evaluate TA.

We demonstrate that TA-Δ_1_ is representative of diverse forms of temporal autocorrelation in our data using a three-fold approach. We restrict our analyses to the HCP, TRT, and Cam-CAN datasets, since they did not use motion scrubbing in their preprocessing pipeline. First, we show that TA-Δ_1_, the first lag term in the ACF, is more reliable than higher lag terms in the ACF. Second, we show that TA-Δ_1_ is predictive of higher lag terms of the ACF, indicating shared information between the terms. Finally, we fit a parametric model of long memory dynamics to our data, and show a strong correlation between the long memory parameter and TA-Δ_1_. These results align with previous work showing TA-Δ_1_ may be representative of higher lag terms of the ACF ^35,104–106^.

Reliability of the ACF decreases with increasing lag. For each lag *k*, we computed the reliability of the corresponding term in the ACF, *ACF*_*x*_(*k*). We found that the median reliability across brain regions decreased as *k* increased, and that TA-Δ_1_ (i.e., *ACF*_*x*_(1)) maximized median reliability (Figure S2a,h). To confirm that this occurs for each brain region individually, we found the difference in reliability between the *ACF*_*x*_(1) and *ACF*_*x*_(*k*), which also decreased across lags (Figure S2b,i). In addition to the reliability of regional TA-Δ_1_, we are interested in how each subject’s mean TA-Δ_1_ (averaging across regions) compares to the mean across other lags. We found that, like in the regional case, reliability decreased as the lag increased (Figure S2c,j). Thus, the reliability at short compared to long time lags motivates a prioritization for TA-Δ_1_ over higher lags.

In addition, TA-Δ_1_ is predictive of higher terms of the ACF. We divided the data into 10 randomly selected subsets for the HCP and Cam-CAN datasets, and 12 subsets for the TRT data. We fit the regression model

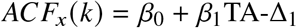

to one of these subsets. Then, we evaluated the *R*^2^ of the model on each of the other subsets. Note that this is a more “difficult” prediction than traditional k-fold cross validation, as it uses less training data to make the predictions. We evaluated this fit on the remaining subsets. Using this procedure, we found that TA-Δ_1_ was predictive of higher lags of the ACF (Figure S2d,k,o). This indicates that information represented in TA-Δ_1_ is partially redundant with that in higher lags.

Furthermore, individual variation in long memory properties of our data can be captured by individual variation in TA-Δ_1_. We focused our analysis on the fractional integration constant *d* from an ARFIMA model, frequently used to capture long memory dynamics^1 107^. Following previous work, we estimated *d* through a univariate wavelet-based Whittle estimator ^108,109^. We found that TA-Δ_1_ is highly correlated with *d* when averaged across subjects (Figure S2e,l,p), averaged across regions (Figure S2f,m,q), or without averaging (Figure S2g,n,r). Under specific assumptions, this relationship between *d* and TA-Δ_1_ can be described analytically ^105,106^. Furthermore, *d* showed similar reliability to TA-Δ_1_ (Figure S2a-c,h-j), and TA-Δ_1_ was predictive of *d* (Figure S2d,k,o). Therefore, in the datasets considered here, TA-Δ_1_ is a reliable and effective measure of TA in general, even though it cannot directly measure long memory dynamics.

## Supplement 2: Mathematical derivations for the spatiotemporal model

### 2.1 Constructing timeseries with specified correlations and power spectra

We seek to create an algorithm that produces timeseries with TA and SA similar to that found in fMRI data. Diverse methods of generating timeseries which exhibit both TA and SA already exist, such as vector autoregressive (VAR) and Ornstein-Uhlenbeck processes. These models directly simulate timeseries in the time domain, introducing TA and SA locally. However, prior research demonstrating powerlaw dynamics in fMRI data ^35,101,102^ suggests that VAR and similar models are inadequate for our purposes, as they are unable to model complex power spectral properties. More complex variants of these models, such as VARFI and FIVAR, are able to capture long memory dynamics, but they are unable to simulate from arbitrary power spectra ^108^. Instead, we seek to generate power spectra which represent the tradeoff between long-memory dynamics, i.e. filtered 1/ *f* ^α^ noise for frequencies above 0.01 hz, and optional white noise, with a flat power spectrum.

To introduce SA in our method, we spatially embed these timeseries with a given covariance matrix *C*. In our case, *C* is given in Equation 2. The usual method of spatially embedding timeseries with a given covariance matrix *C* is to multiply them by the matrix square root of *C*, equivalent to sampling from a multivariate normal distribution. However, this process involves the linear combination of distinct timeseries, which changes their power spectra, destroying the desired power spectrum. To avoid this confound, we introduce the following algorithm, a generalization of Ref. ^110^, which is able to produce correlated timeseries with arbitrary power spectra. The algorithm can alternatively be seen as a generalization of phase randomization to obtain specified correlational structure.

**Algorithm** Let *N* be the number of desired timeseries, and let *T* be the even length of the desired timeseries. Given an *N* × *N* correlation matrix *C*_*i, j*_ and a set of exponents *α*_*i*_, we seek to generate a set of timeseries {*x*_1_[*t*], …, *x*_*N*_ [*t*]} such that each timeseries *x*_*i*_[*t*] has a specified estimate of the power spectrum | *X*_*i*_[*k*]|^2^ and the correlation matrix of the resulting timeseries has expected value *C*_*i, j*_. The following algorithm operates by generating a set of complex frequency-domain coefficients in the domain [−*T* /2 + 1, *T* /2], introducing SA and TA independently, and then performing an inverse discrete Fourier transform to create the desired set of timeseries.

1. Pick the desired power spectra estimates {| *X*_1_[*k*]|^2^, …, | *X*_*N*_ [*k*]|^2^}, representing the desired temporal structure, and *N* × *N* correlation matrix *C*, representing the desired spatial structure. These choices must satisfy the requirement that the matrix

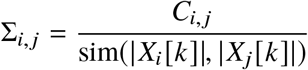

is positive semi-definite, where sim denotes cosine similarity, i.e.,

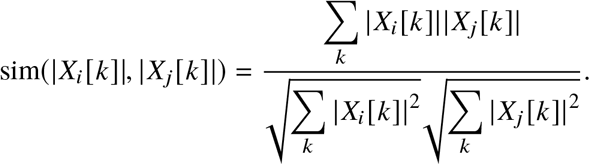
2. For each region *n* ∈ {1, …, *N*}, and for each frequency *k* in the positive sub-Nyquist Fourier domain, 1 ≤ *k* ≤ *T* /2, sample two vectors from a multivariate normal distribution 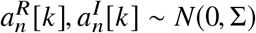.
3. Set 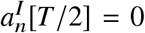 for all *n*. Because *T* is even, by way of symmetry the frequency-domain coefficients of the Nyquist frequency must be real. Set 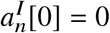 so that the resulting timeseries will be real.
4. Form the complex frequency-domain coefficients by letting 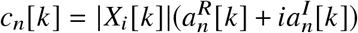 for 0 ≤ *k* ≤ *T* /2, and 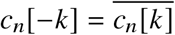 for 0 *< k < T* /2.
5. Perform an inverse discrete Fourier transform for each *c*_*n*_[*k*] to obtain the timeseries *x*_*n*_[*t*].

Each of the resulting timeseries *x*_*i*_[*t*] will have a frequency spectrum approximately equal to 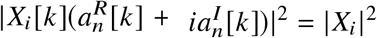, the desired temporal structure. To show that {*x*_1_[*t*], …, *x*_*N*_ [*t*]} have the desired spatial structure, we show that for any *i, j*, 𝔼 [corr(*x*_*i*_[*t*], *x*_*j*_[*t*])] = *C*_*i, j*_. As an intermediate step, we show that 𝔼 [corr(*x*_*i*_[*t*], *x*_*j*_[*t*])] = corr(Re(*X*_*i*_[*k*]), Re(*X*_*j*_[*k*]))).

Let *x*_*i*_[*t*] and *x*_*j*_[*t*] be two of the timeseries resulting from this algorithm. Note that, for large *N*, corr(*x*_*i*_[*t*], *x*_*j*_[*t*]) = cov(*x*_*i*_[*t*], *x*_*j*_[*t*])/(var(*x*_*i*_[*t*]) · var(*x*_*j*_[*t*]))^1/2^, where var and cov are the empirical estimates of the variance and covariance, respectively. We first seek to expand cov(*x*_*i*_[*t*], *x*_*j*_[*t*]), var(*x*_*i*_[*t*]), and var(*x*_*j*_[*t*]). Without loss of generality, we may assume that *x*_*i*_[*t*] and *x*_*j*_[*t*] have a mean of 0, as their mean does not influence the spatial or temporal structure of the set of timeseries, so we can estimate cov(*x*_*i*_[*t*], *x*_*j*_[*t*]) = 1/*N Σ*_*t*_ *x*_*i*_[*t*]*x*_*j*_[*t*].

Recall that *X*_*i*_[*k*] is the discrete Fourier transform ℱ (*x*_*i*_[*t*]). We denote the even and odd components 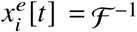 and 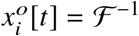. Thus, our estimator of the covariance can be written as

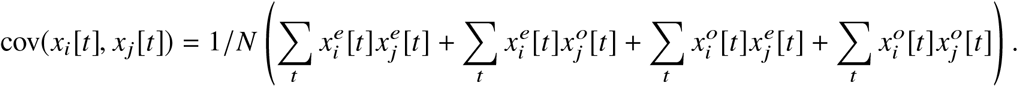

For the even part of the function, 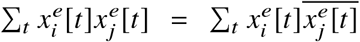, and by Parseval’s theorem, this equals 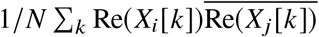. This can be written as 1 / *N Σ*_*k*_ Re (*X*_*i*_ [*k*]) Re (*X*_*j*_ [*k*] = cov(*X*_*i*_ [*k*] Re (*X*_*j*_ [*k*]). By the some logic, 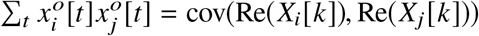.

The real and imaginary components of *X*_*i*_[*k*] are independent by construction, which means 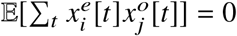 and 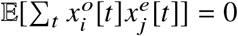. Thus, we can rewrite the covariance estimator as

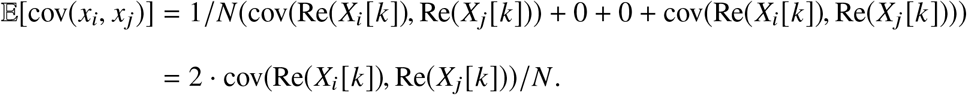

This gives 𝔼 [var(*x*_*i*_)] = 𝔼 [cov(*x*_*i*_, *x*_*i*_)] = 2 · var(Re(*X*_*i*_[*k*]))/*N*, so

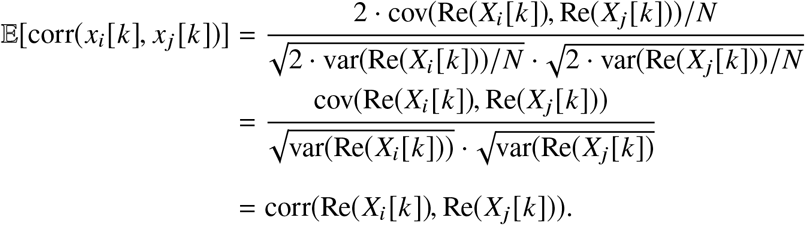

A similar procedure applies to the imaginary component of *X*_*i*_[*k*].

We now show that 𝔼 [corr(Re(*X*_*i*_[*k*]), Re(*X*_*j*_[*k*]))] = *C*_*i, j*_. We note that, by construction, 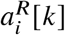 and |*X*_*i*_[*k*]|^2^ are independent, and that 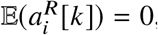, meaning 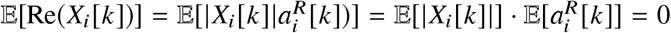. This allows us to expand as before, giving

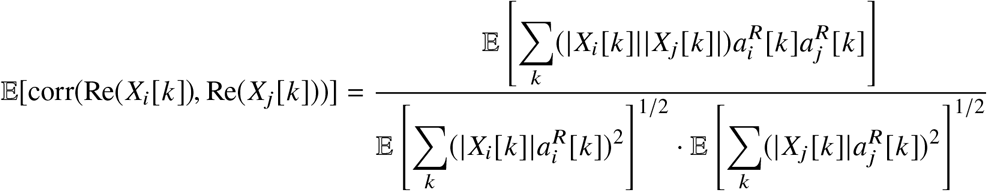

Because 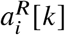 and 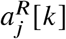 have mean 0 and variance 1 and have correlation equal to approximately Σ_*i, j*_, the product 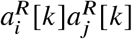 has an expected value of Σ_*i, j*_. By construction, 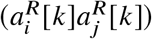 and (|*X*_*i*_[*k*]|| *X*_*j*_[*k*]|) are independent, so

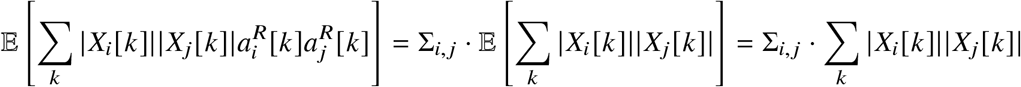

When *i* = *j, C*_*i, j*_ = 1. By the same logic,

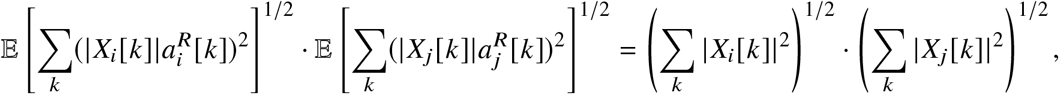

and so

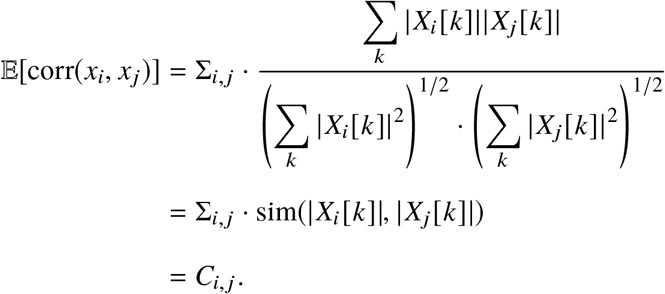

### 2.2 Relationship between TA-Δ_1_ and power spectrum

Our primary measure of temporal autocorrelation is TA-Δ_1_, despite the fact that our model uses timeseries with complicated power spectra. To demonstrate that TA-Δ_1_ can capture important information about the power spectrum, we derive the relationship between the Fourier spectrum and TA-Δ_1_.

Suppose we have a finite timeseries *x*[*t*] with discrete Fourier transform *X*[*k*]. We claim that its TA-Δ_1_ is

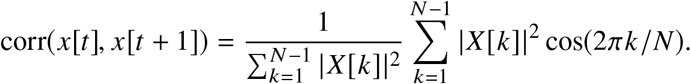

Let *ACF*(*x*)[*τ*] denote the autocovariance of *X* at lag *τ*, i.e. cov(*x*[*t*], *x*[*t* + *τ*]). By the Wiener-Khinchin theorem, *ACF*(*x*)[*τ*] is the inverse Fourier transform of an estimate of the power spectrum, | *X*[*k*]|^2^. Without loss of generality, let us suppose *x*[*t*] has zero mean, and hence, *X*[0] = 0, so that

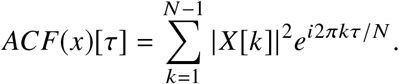

Because *x*[*t*] is real, *e*^*i*2π*k*τ/*N*^ = cos(2*πkτ*/*N*). Making the approximation that var(*x*[*t*]) = var(*x*[*t* + 1]), we have

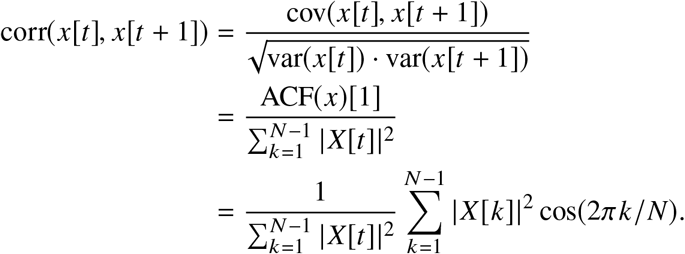

### 2.3 Relationship between TA-Δ_1_ and noisy power spectrum

Additionally, we would like to know how the addition of white noise to a power spectrum will change the TA-Δ_1_. We prove that, for an finite timeseries *y*[*t*] = *x*[*t*] + *w*[*t*] where *w*[*t*] ∼ *N*(0, *σ*) of length *N*,

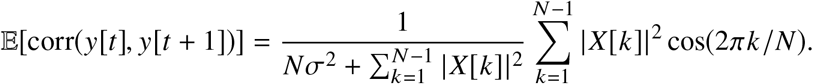

The Fourier transform is linear, so we can write

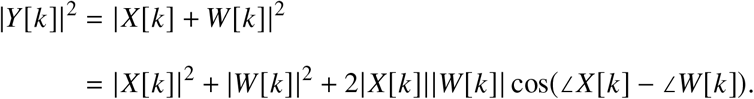

Since *w*[*t*] is white noise, *W* [*k*] has the properties 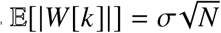 and ∠*W* [*k*] ∼ *U*(0, 2*π*). Thus, 𝔼 [cos(∠*X*[*t*] − ∠*W* [*t*])] = 0, so

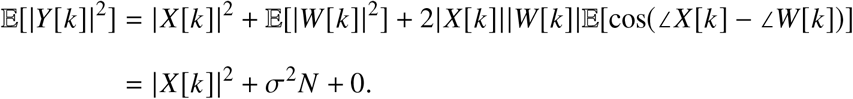

Applying our result from Supplement 2.2, we see that

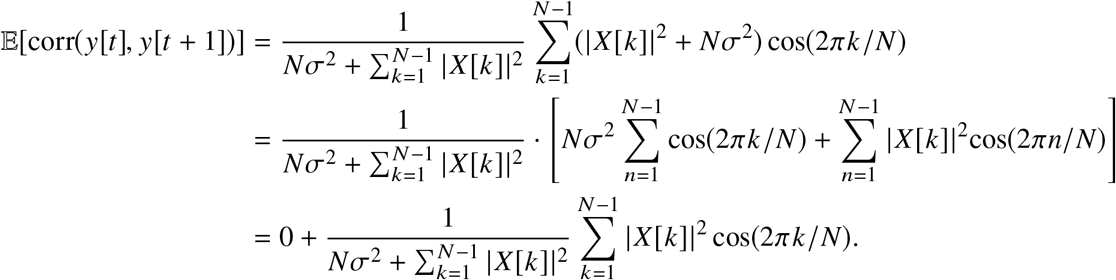

It directly follows that, for a given 𝔼 [corr(*y*[*t*], *y*[*t* + 1])] = *ϕ*, we can solve this equation for *σ*^2^ as

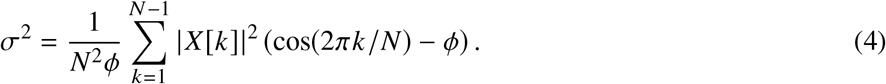

### 2.4 Parameter-free fitting of the spatiotemporal model

In theory, it is possible to fit the spatiotemporal model in a parameter-free manner. In other words, the SA-λ, SA-∞, and TA-Δ_1_ statistics can all be measured from the data, and therefore, it should be possible to choose SA-λ^gen^ and SA-∞^gen^, with arbitrary power spectra estimators | *X*_*i*_[*k*]|^2^ for region *i*, such that the spatiotemporal model produces timeseries which have a given SA-λ, SA-∞, and TA-Δ_1_. We describe the mathematical derivation of a procedure for selecting SA-λ and SA-∞ using this logic, and show why this procedure is infeasible in practice.

As a reminder, the spatiotemporal model has four steps: (1) use SA-λ^gen^ and SA-∞^gen^ in conjunction with the nodal Euclidean distances *d*_*i, j*_ to generate a spatial correlation matrix *C*; (2) construct a power spectrum corresponding to high-pass filtered Brownian (1/ *f* ^2^) noise; (3) construct timeseries *x*_1_, …, *x*_*N*_ for brain regions 1, …, *N* according to Supplement 2.1 such that corr(*x*_*i*_, *x*_*j*_) = *C*_*i, j*_ and all timeseries have the power spectrum specified in step (2); (4) for each time point *t* in each timeseries *x*_*i*_[*t*], add white noise 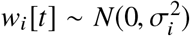, with 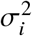 chosen according to Supplement 2.3 such that *x*_*i*_[*t*] has TA-Δ_1_ constant TA-Δ_1*i*_.

To perform the derivation, suppose that we have two timeseries *x*_*i*_ and *x*_*j*_, *i* ≠ *j*, such that 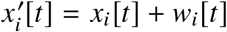, where *x*_*i*_[*t*] has some power spectrum | *X*_*i*_[*t*]|^2^ for all *i* and 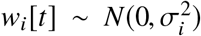 for given 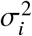. Let the correlation corr(*x*_*i*_, *x*_*j*_) = *C*_*i, j*_ for some unknown *C*_*i, j*_. We derive an equation to select *C*_*i, j*_ such that 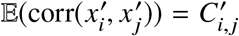 for some desired 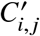

To begin, we rewrite

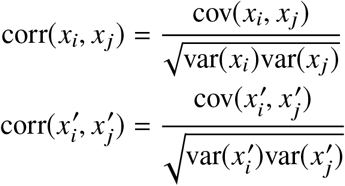

Since 𝔼 (cov(*x*_*i*_ + *w*_*i*_, *x*_*j*_ + *w*_*j*_)) = cov(*x*_*i*_, *x*_*j*_) for *i* ≠ *j*, and 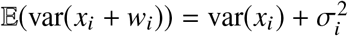, we can write

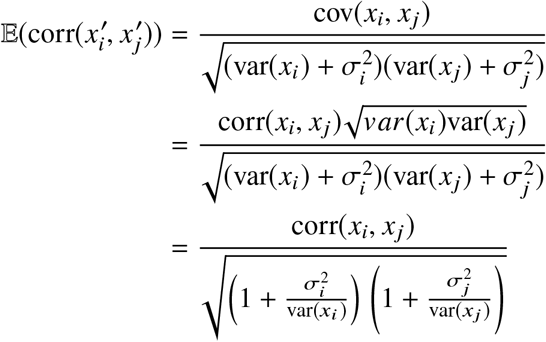

The variance of a timeseries *x*_*i*_ can be estimated by its power spectrum using Parseval’s theorem, such that 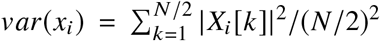. Thus, substituting 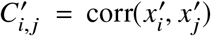 and *C*_*i, j*_ = corr(*x*_*i*_, *x*_*j*_) and rearranging terms, we have

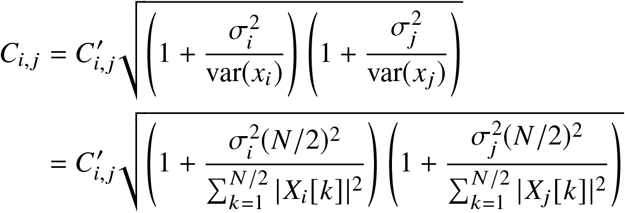

Thus, we have derived a *C*_*i, j*_ which can be used in our procedure to produce timeseries with correlation 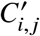.

While mathematically interesting, the derivation is not useful in practice. Because we are computing a correlation, *C*_*i, j*_ has an upper bound of 1, thereby bounding the potential values of 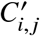 which can be produced. In practice this constraint is difficult to satisfy. In our data, 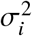 can take on values up to an order of magnitude larger than var(*x*_*i*_), meaning that only very small values of 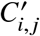 (approximately 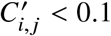) can satisfy the constraint. Additionally, in the analysis of neural data, we consider more than two timeseries. This means that *C* must be a correlation matrix, and thus *C* must be positive semi-definite, or equivalently, have all non-negative eigenvalues. In our data, approximately 25% of eigenvalues are negative after applying this procedure, showing that it is unlikely to be useful in practice.

### Supplement 3: Relationship between temporal autocorrelation and variance in correlation

Previous work established that temporal autocorrelation (TA) increases the variance of the Pearson correlation between pairs of timeseries ^31,34,111,113^. Here, we quantified TA using TA-Δ_1_, the TA at a single lag. We did this with the understanding that TA-Δ_1_ correlates with TA at higher lags, or with an underlying long memory process (Supplement 1). Since higher lag TA may have an impact on the variance, it is insufficient to solely measure the effect of TA-Δ_1_ on the variance in Pearson correlation. A more general approach is to quantify the effect of the entire power spectrum on the variance in Pearson correlation (Supplement 2.2). The power spectrum is a full-rank linear transformation of the autocorrelation function, and therefore, is a non-parametric analysis of TA for both short and long memory dynamics. Thus, we seek to understand the impact of an arbitrary power spectrum on the variance in Pearson correlation of two timeseries.

We focus on a specific case of particular importance by analyzing two independent timeseries with identical power spectra and phases distributed randomly around the unit circle. We show that the variance of the Pearson correlation between these two timeseries increases as the uniformity of the power spectrum decreases. In other words, Pearson correlation is lowest for a timeseries of iid white noise, which by definition has no TA. It increases as the power spectrum deviates from uniformity, and is highest for spectra consisting of a single frequency, i.e. a sinusoid.

The variance of a timeseries is closely connected with the power spectrum. Consider the real, discrete, finite timeseries *x*_1_[*t*] and *x*_2_[*t*] of length *N*, with Fourier transform *X*_1_[*k*] and *X*_2_[*k*] and variance 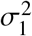 and 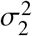. Without loss of generality, assume *x*_1_[*t*] and *x*_2_[*t*] have mean 0. We can estimate the covariance for large *N* as

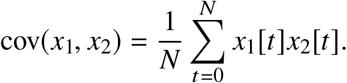

Then, by Parseval’s theorem, we can write the covariance as

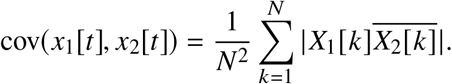

Therefore, by noting that the variance of a timeseries is simply self-covariance, the empirical Pearson correlation corr can be written as

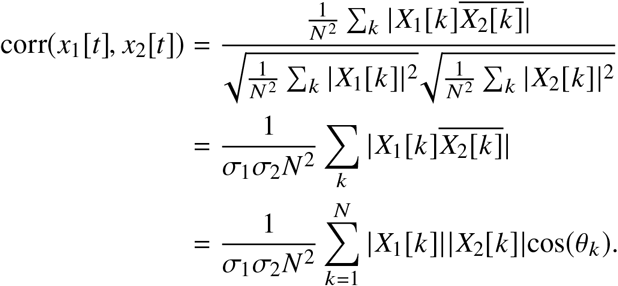

Note that the quantities | *X*_1_[*k*]|^2^ and | *X*_2_[*k*]|^2^ are estimates of the power spectra of *x*_1_[*t*] and *x*_2_[*t*]. Since we assume the angles *θ*_*k*_ are selected randomly from the uniform distribution from [0, 2*π*], we can easily compute

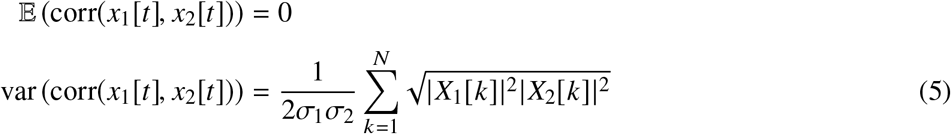

The variance given in Equation 5 is maximized at 1/2*σ*_1_*σ*_2_ when both power spectra have unit spikes at the same location, corresponding to sinusoids with identical frequency. Since we assume that *x*_1_[*t*] and *x*_2_[*t*] have identical power spectra, i.e. | *X*_1_[*k*]|^2^ = | *X*_2_[*k*]|^2^, Equation 5 is minimized at 1/2*σ*_1_*σ*_2_*N* when *x*_1_[*t*] and *x*_2_[*t*] are white noise, i.e. | *X*_1_[*k*]| = 1/*N*. (If we had not assumed the power spectra were identical, the variance would be minimized at zero for any non-overlapping power spectra, indicating that it is impossible for these timeseries to show a non-zero Pearson correlation.) Hence, any deviation from a uniformly-distributed power spectrum, as occurs in temporally-autocorrelated timeseries, will increase the variance in the Pearson correlation.

## Supplement 4: Economic clustering model

### 4.1 Overview

Our analysis focused on models at the level of time series. However, another popular type of generative model simulates network formation by directly adding edges to a set of nodes. We refer to these models as “generative graph models” to distinguish them from timeseries-based generative models. Generative graph models have been especially useful in modeling real-world networks ^115,116^. For example, a generative graph model can capture the property that nodes which are already well-connected are likely to gain even more connections, i.e., that the “rich get richer” ^3^. It effectively models the generative process of real-world networks such as social networks, where people with a large number of friends are more likely to gain new friends over time. By simulating how real networks are formed, generative graph models reveal the mechanisms that drive the high-level structure of those networks. For this reason, these models have quickly become ubiquitous in the natural and social sciences.

Brain networks share many properties with other complex biological and physical systems, so they can be analyzed using similar graph-theoretic methods. Recently, generative graph models have been used in neuroscience to model the human connectome ^5,119,120^, opening up new avenues of analysis of both functional ^22,24^ and structural ^23,123,124,126,127^ connectivity data. One reason why generative graph models are particularly attractive in the field is that they appear, at first glance, to model the trade-off between wiring cost and topological efficiency in brain networks ^128^. In particular, generative graph models that construct networks based on the physical distance between brain regions and a few basic topological properties of the network have been especially effective ^22,23^. In addition, generative graph models are relatively simple to construct and conceptualize, and they can be analyzed using computational tools from network science and graph theory.

Despite their popularity and conceptual simplicity, most generative graph models are removed from the underlying biological processes that occur in the brain. This is because the generative process used to construct those models is starkly different from the actual pipeline used to extract connectivity data from brain signals. Nevertheless, these models, such as the economical clustering (EC) model ^22^, are exceedingly effective at explaining the topology of brain connectivity.

In order to explain this effectiveness using a more biologically-motivated timeseries-based generative process, we compare the EC model from Ref. ^22^ directly to our spatiotemporal model. The EC model is controlled by two parameters: one controls the connection probability with respect to distance, and the other with respect to homophily (i.e. shared neighbors, or clustering) between nodes ^22^. There are a few apparent symmetries between the EC model and our spatiotemporal model. Both models produce realistic synthetic networks, and they are each driven by two features, one of which relates connectivity with physical distance. These initial similarities suggest the possibility that the second parameter in each model may also capture similar underlying properties. In fact, our findings show that both pairs of parameters in the two models are indeed tightly correlated, which opens up a new biologically-motivated interpretation of the generative graph model.

### 4.2 Economic clustering model

We compared the spatiotemporal model to the “economic clustering” (EC) model ^22^, which was chosen because of its good performance and its apparent parallels to our timeseries-based model. We briefly summarize the model here. We constructed a minimum spanning tree of the mean FC matrix using Kruskal’s algorithm ^129^ to ensure that our procedure does not produce disconnected components. Then, edges were added one at a time according to a probabilistic wiring rule, until the number of edges in the model network and observed network were equal. The relative probability of adding a connection between node *u* and node *v* was determined by:

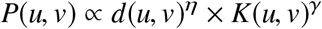

where *d*(*u, v*) is the Euclidean distance between nodes *u* and *v, K*(*u, v*) is the number of shared neighbors between nodes *u* and *v*, described below, and *η* (the distance parameter) and *γ* (the clustering parameter) are free parameters. The exponent *η* controls the extent to which distance impacts connection probability. For highly negative *η*, shortrange connections are much more likely; for values of *η* close to 0, all connections regardless of distance are almost equally likely to be added.

*K*(*u, v*) represents the number of shared neighbors between nodes *u* and *v*, or equivalently, the number of nodes that are connected to both node *u* and node *v*. We can compute this as *K*(*u, v*) = Σ_*w*_ *A*_*uw*_ *A*_*wv*_ where *A* is the adjacency matrix of the graph, i.e. *A*_*ij*_ = 1 if nodes *i* and *j* are connected, and *A*_*ij*_ = 0 otherwise. Note that *K*(*u, v*) must be recomputed at each iteration of the algorithm, since the number of shared neighbors can change after an edge is added. The exponent *γ* controls the extent to which the number of shared neighbors impacts connection probability. For *γ <* 0, nodes with few shared neighbors are more likely to be connected; for *γ >* 0, nodes with many shared neighbors are more likely to be connected.

Note that this procedure does not generate networks which vary smoothly with the parameters *η* and *γ*. In other words, for a given seed, a small change in either of these parameters could cause major changes in the topology of the generated graph. This occurs because each edge addition causes a cascading change in connection probabilities, and thus, even slight differences in the first edges added will have a large impact on the generated graph.

It has been shown that generative graph models which only specify connection probability as a function of distance do not capture features such as small-worldness, modularity, and degree distribution of real brain networks; rather, a second parameter is necessary to emulate these topological properties ^22^. These findings are consistent with the results of our spatiotemporal model. In particular, our spatiotemporal model has two main distinguishing features: spatial autocorrelation and temporal autocorrelation. This apparent symmetry between the spatiotemporal model and the EC model motivates our comparison of the underlying parameters across the two models.

### 4.3 Fitting the spatiotemporal model to the EC model

To determine whether the parameters of the spatiotemporal model and the EC model are correlated, we developed a fitting procedure linking the two models. Because the model is determined by probabilistic wiring rules, a small perturbation in a model parameter may cause a large change in graph structure. Thus, the highly stochastic nature of the EC model produces a rugged optimization landscape, which prevents us from fitting the parameters *η* and *γ* using conventional optimization algorithms. In order to circumvent this problem, we instead fit the spatiotemporal model to instances of the EC model, since the spatiotemporal model has a smooth optimization landscape.

Overall, the procedure consists of the following steps. For a given pair of (*η, γ*), we generate ten instances of the EC model. Then, we fit the spatiotemporal model to the population of simulated EC models and record the optimal parameters SA-λ and TA-Δ_1_. We repeat these steps over a predetermined range of *η* and *γ* values that generally produce reasonable networks. On a high level, this procedure returns the parameters SA-λ and TA-Δ_1_ of the spatiotemporal model that fit best to the EC model determined by given values of *η* and *γ*. This allows us to conduct further analyses to reveal how changes in the EC model parameters affect changes in the corresponding spatiotemporal model parameters, thereby linking the two models for comparison.

In order to perform this fitting procedure, we need to establish a similarity metric between the multiple instances of the EC model and the single instance of the spatiotemporal model. We also require an optimization method for fitting parameters that maximize this similarity metric.

To evaluate the similarity between two networks, we use the energy function presented in Ref. ^23^,

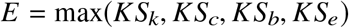

where each argument in the max function is the two-sample Kolmogorov-Smirnov test statistic between the networks’ degree (*k*), clustering (*c*), betweenness centrality (*b*), and edge length (*e*) distributions. Note that for consistency with Ref. ^23^, this energy function measures fitness based on nodal graph metrics, in contrast to the energy function for fitting the spatiotemporal model to data, which is based on eigenvalues.

To reduce noise and to produce population-level fits, we collectively minimize the average energy between all instances of the EC model and a single instance of the spatiotemporal model,

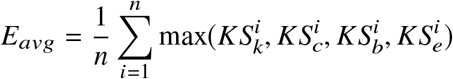

where *n* is the number of subjects in our dataset, and each argument to the max function is the Kolmogorov-Smirnov statistic between the distributions of *i*th instance of the generative graph model and the spatiotemporal model for the corresponding graph metric. Instances of the EC model differ only in the initial random seed. We used differential evolution to determine the optimal model parameters.

### 4.4 Results

One apparent similarity between the construction of the EC model and the homogeneous spatiotemporal model is that each model includes a parameter that controls connection probability with respect to distance between nodes: the distance parameter *η* in the EC model and the SA-λ parameter in the homogeneous spatiotemporal model. As *η* → 0^−^, distance plays a smaller role in determining connection probability, so long- and short-range connections are more equally likely to be chosen. As SA-λ → 0^+^, the spatial autocorrelation approaches zero for all non-zero distances, so distance has a diminishing effect on the correlation. Therefore, we expect the EC distance parameter *η* to be negatively correlated with SA-λ. To test this empirically, we fix the EC clustering parameter at several different values and vary the EC distance parameter, fitting the homogeneous spatiotemporal model to each of the EC model’s simulated networks. We find a strong negative relationship between the EC distance parameter *η* and SA-λ (Figure S8a). This confirms the similarities expected between the two spatial parameters.

The ability of both models to capture network topology, combined with the strong association of SA-λ and EC distance parameter, suggests that the second parameters of these two models may also be related. Specifically, we test whether the clustering parameter *γ* in the EC model and TA-Δ_1_ in the homogeneous spatiotemporal model are associated as well. After applying the same procedure of simulating from the EC model and fitting with the homogeneous spatiotemporal model, we found a tight relationship between TA-Δ_1_ and the EC clustering parameter (Figure S8b). Indeed, varying these parameters in the two models results in similar changes in graph metrics (Figure S8). By contrast, the TA-Δ_1_ parameter is not strongly associated with the EC distance parameter (Figure S8c), and the SA-λ parameter is not strongly associated with the EC clustering parameter (Figure S8d). Thus, increases in SA breadth reduce the relative probability of nearby connections, and increases in TA increase the probability of connections occurring preferentially within clusters. This confirms our intuition of how SA and TA influence graph topology.

---

**Figure S1.**
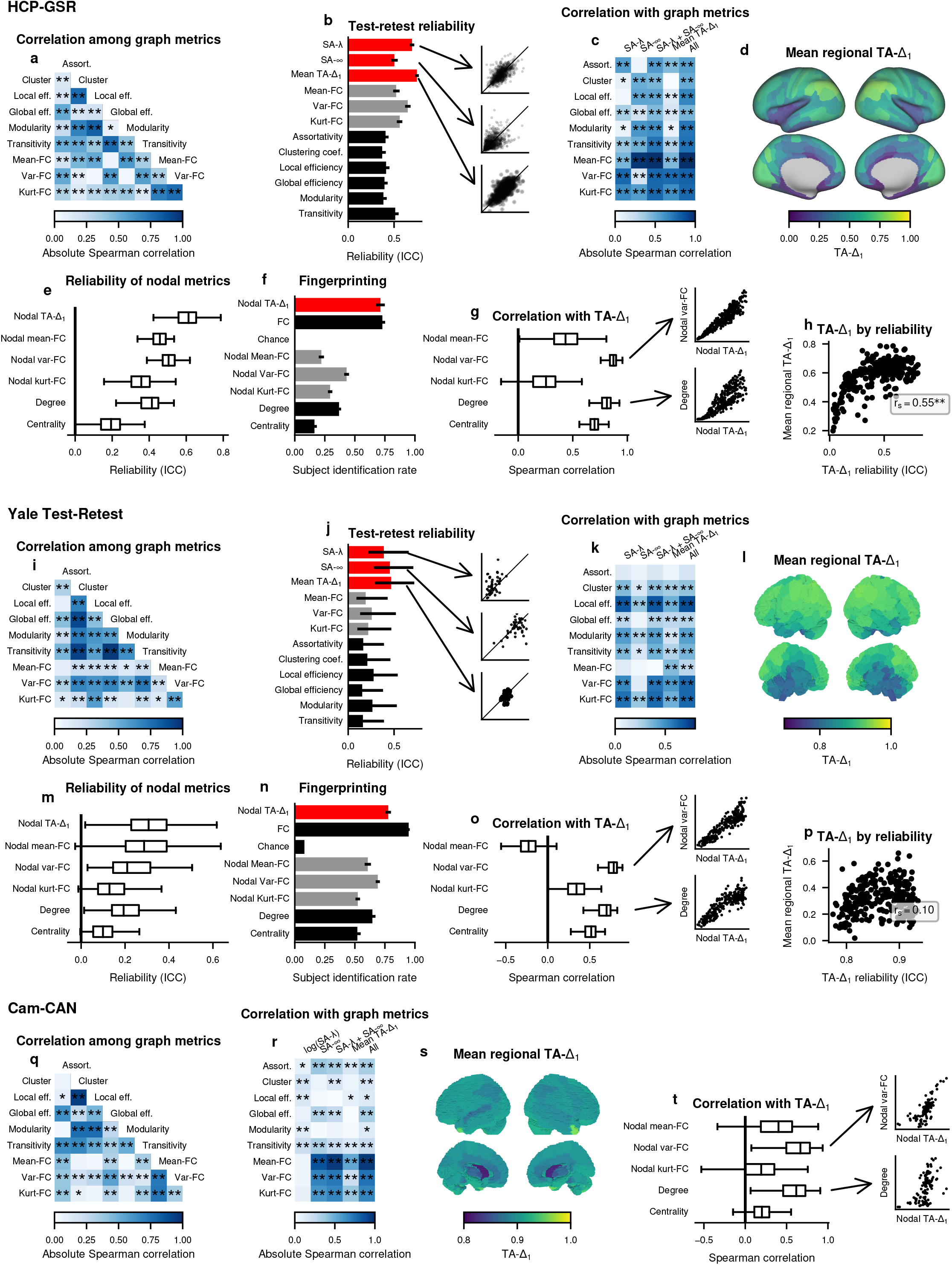
SA and TA are important features of rs-fMRI. Similar format as Figure 1, but for the HCP-GSR, TRT, and Cam-CAN datasets.

**Figure S2.**
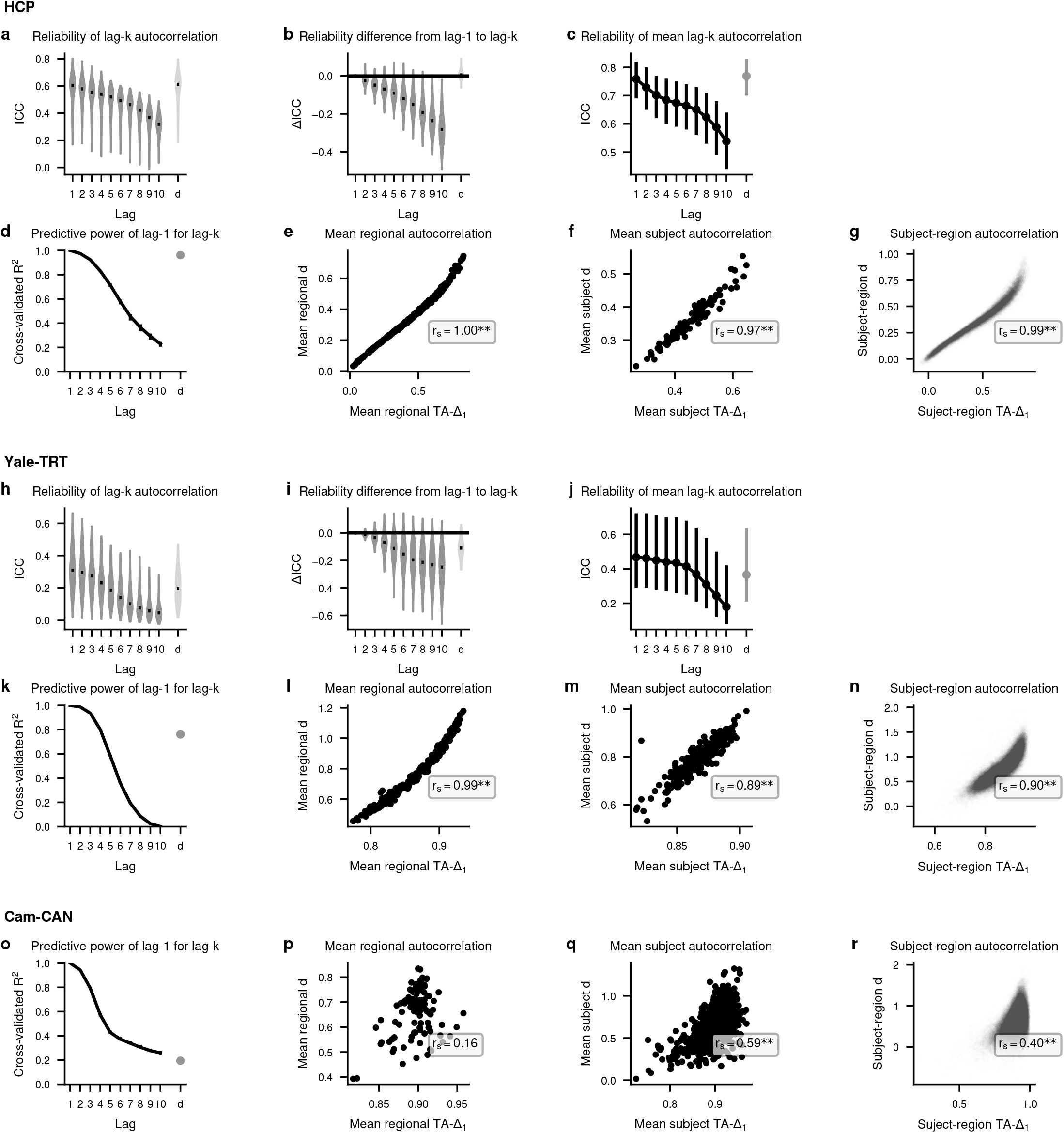
TA-Δ_1_ captures individual variation in long memory dynamics. (a,h) For each brain region, intraclass correlation coefficient (ICC) was computed across runs for the same subject for various lags of the temporal autocorrelation function (ACF). The distribution of reliability across regions is shown in gray, with the median in black. *d* indicates the fractional integration constant from an ARFIMA(0,d,0) model fit to the timeseries. (b,i) The distribution of differences in reliability (ΔICC) between each lag in the ACF and TA-Δ_1_. *d* indicates the difference in reliability between TA-Δ_1_ and the fractional integration constant. (c,j) The reliability of the mean TA-Δ_1_ for each lag, averaged across all regions in each subject, where error bars are 95% confidence intervals. *d* indicates reliability of the mean fractional integration constant. (d,k,o) A linear model predicted each term of the ACF using TA-Δ_1_. The model was fit to one subset of the data, and tested on the remaining subsets. Error bars indicate the maximum and minimum *R*^2^ among the subsets, and are sometimes hidden beneath the curve. *d* indicates predictions of the fractional integration constant. (e-g,l-n,p-r) Estimates of TA-Δ_1_ are compared to estimates of *d*. Values were computed for each region, averaging across subjects (e,l,p); for each subject, averaging across regions (f,m,q); and for each subject and region, without any averaging (g,n,r). *r*_*s*_ indicates Spearman correlation, where * indicates p<.05, and ** indicates p<.01.

**Figure S3.**
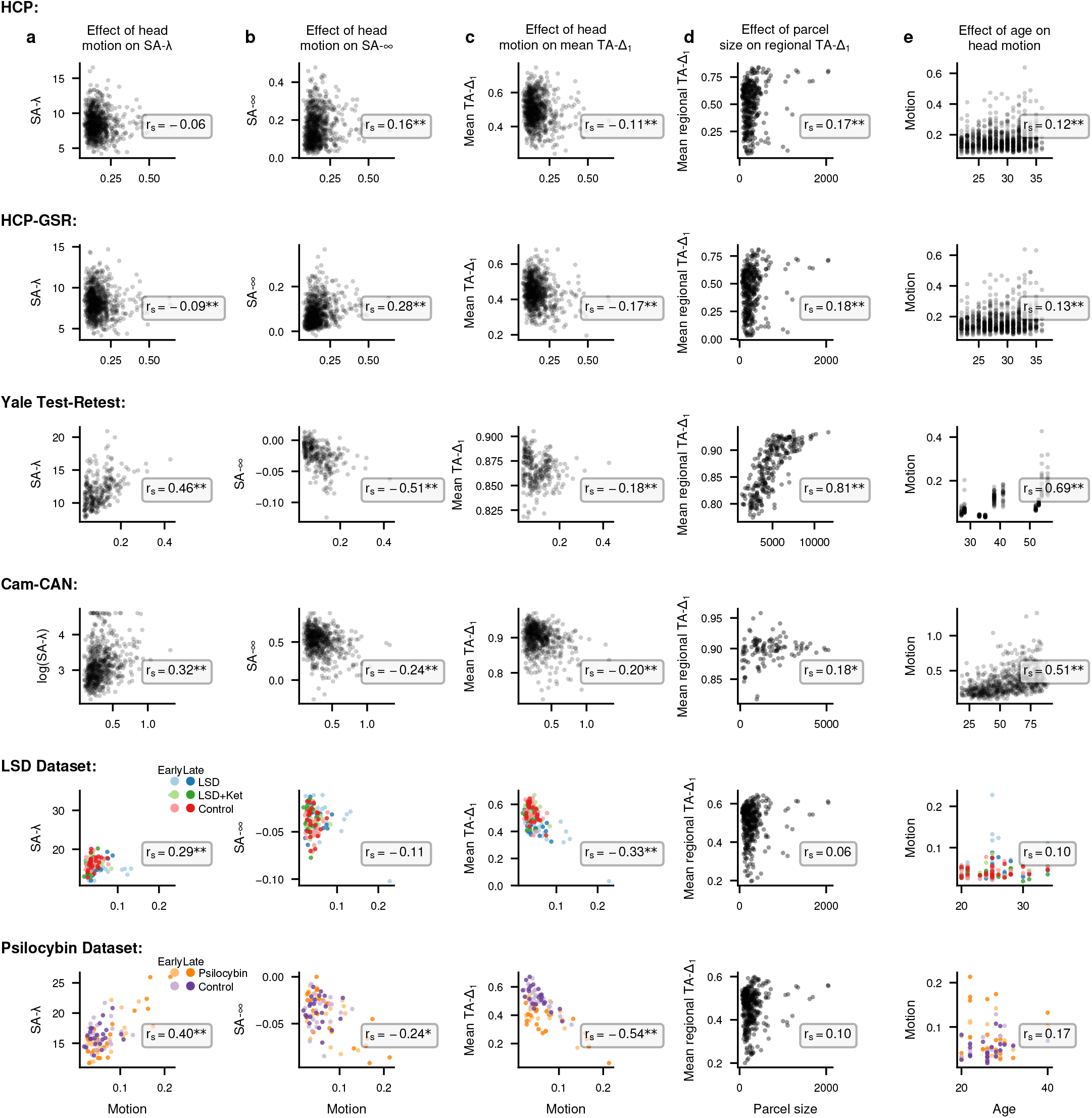
Correlation of motion and parcel size with spatial and temporal autocorrelation. (a-c) For each dataset, the mean framewise displacement (“motion”) is compared to (a) SA-λ, (b) SA- ∞, and (c) mean TA-Δ_1_, with inset Spearman correlation (*r*_*s*_). (d) The average TA-Δ_1_ for each parcel across all subjects is plotted against the parcel’s size, with inset Spearman correlation. Parcel size is measured in surface area for HCP and HCP-GSR, and number of voxels for Yale Test-Retest and Cam-CAN. (e) Subject’s mean framewise displacement (“motion”) is plotted against the subject’s age. * indicates Spearman correlation p<.05, and ** indicates p<.01.

**Figure S4.**
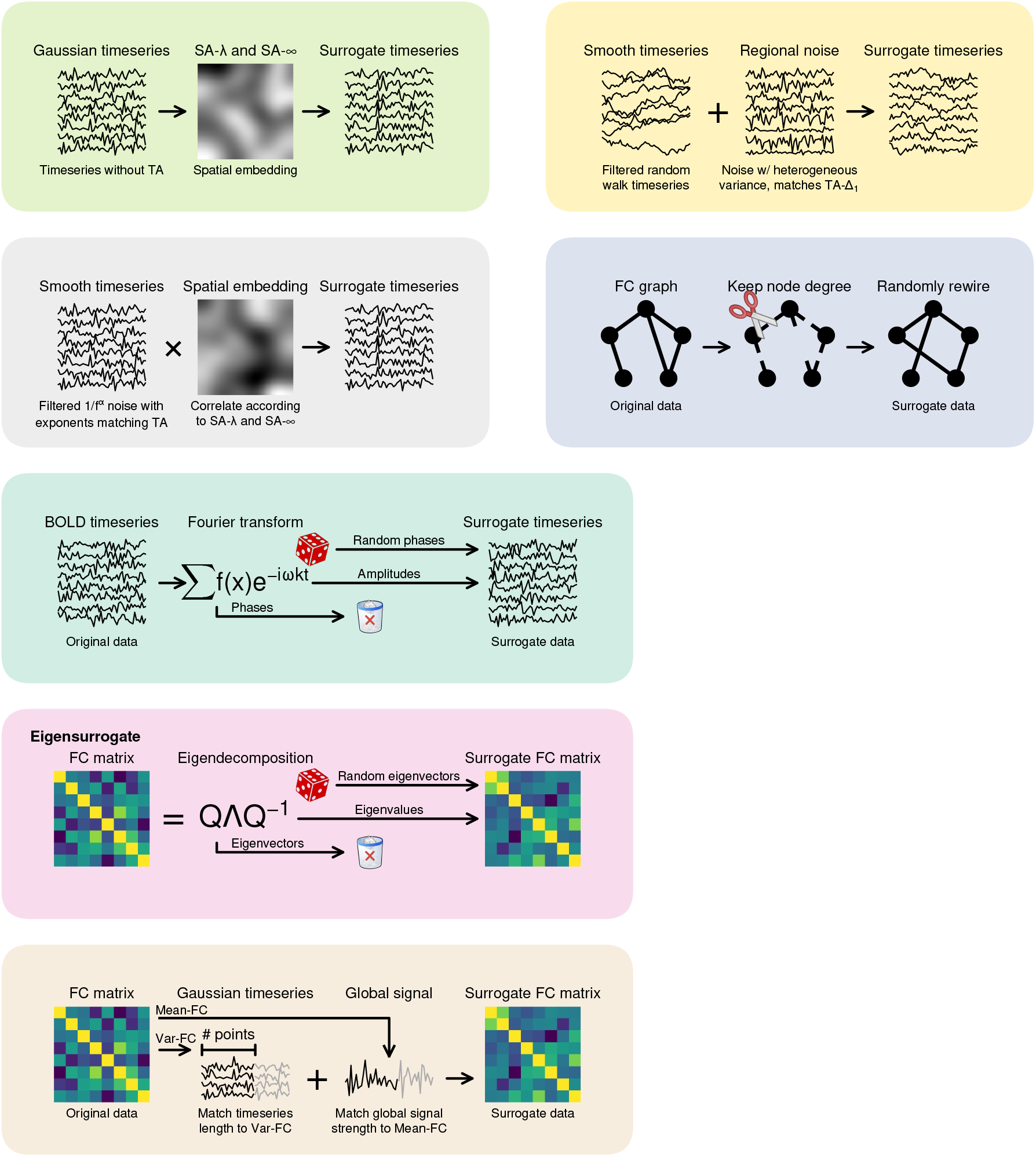
Schematics of all of the models. All models considered are described in their corresponding schematics.

**Figure S5.**
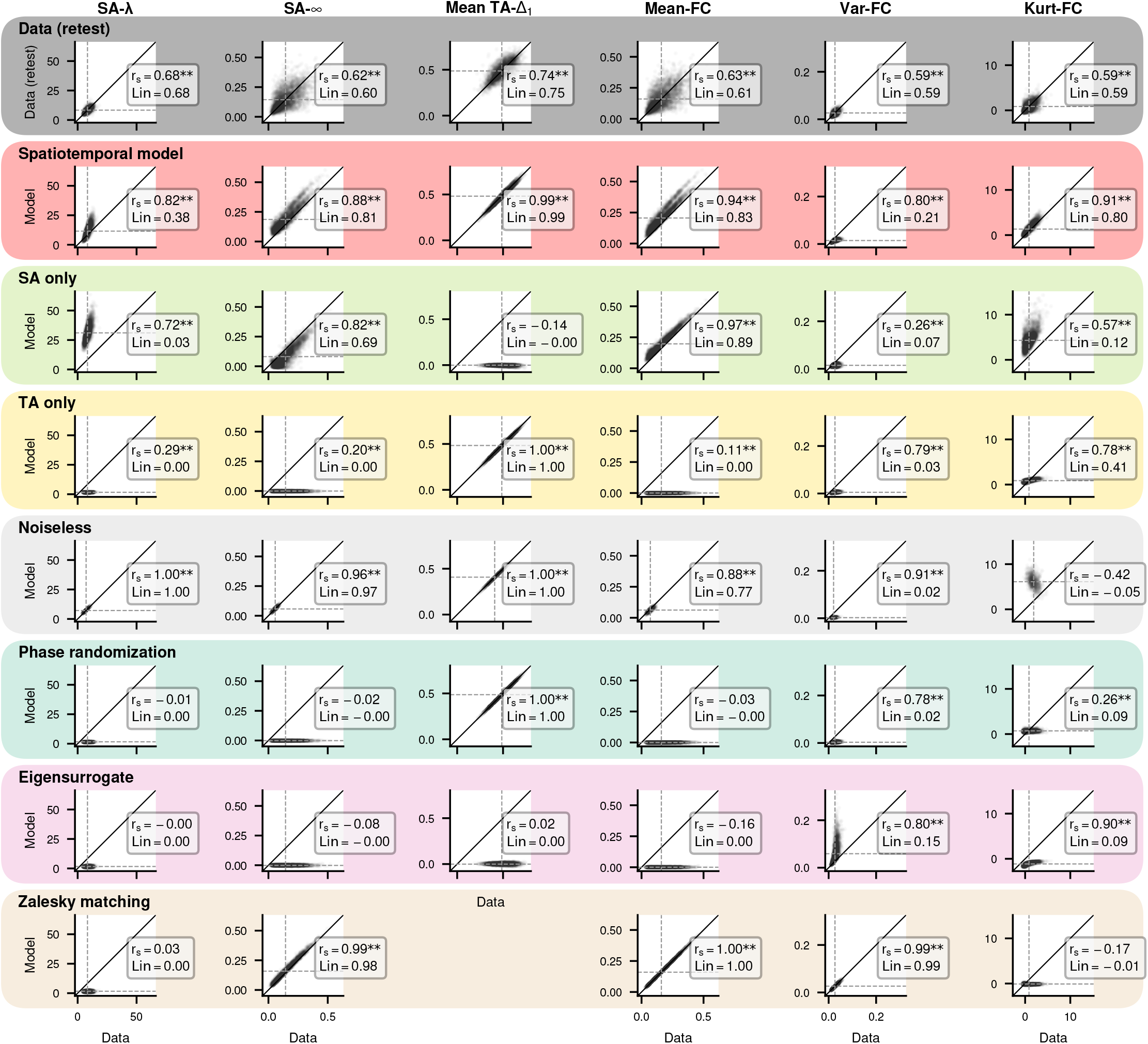
Correlation of model and data graph metrics for all models in HCP data. For each model, each subject’s empirical graph metrics are plotted against model graph metrics for metrics from Figure 2i-p for the HCP dataset. Spearman correlation (*r*_*s*_) and Lin’s concordance (Lin) are inset. * indicates Spearman correlation p<.05, and ** indicates p<.01. Zalesky matching operates at the level of the FC matrix, and thus, TA-Δ_1_ could not be computed. Edge reshuffle operates at the level of the graph, preventing any of these measures from being computed.

**Figure S6.**
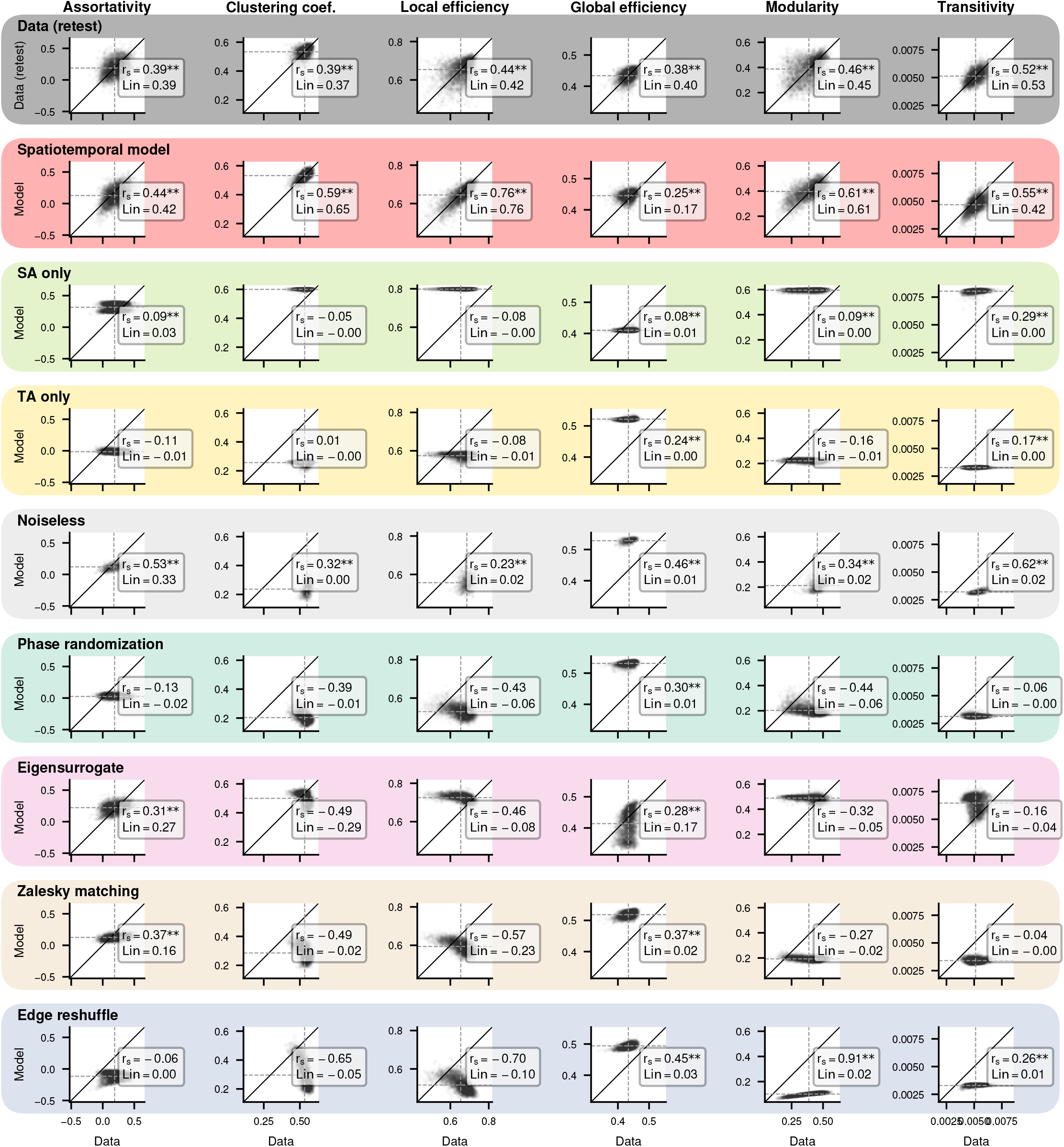
Correlation of model and data graph metrics for all models in HCP data. For each model, each subject’s empirical graph metrics are plotted against model graph metrics for metrics from Figure 2i-p for the HCP dataset. Spearman correlation (*r*_*s*_) and Lin’s concordance (Lin) are inset. * indicates Spearman correlation p<.05, and ** indicates p<.01.

**Figure S7.**
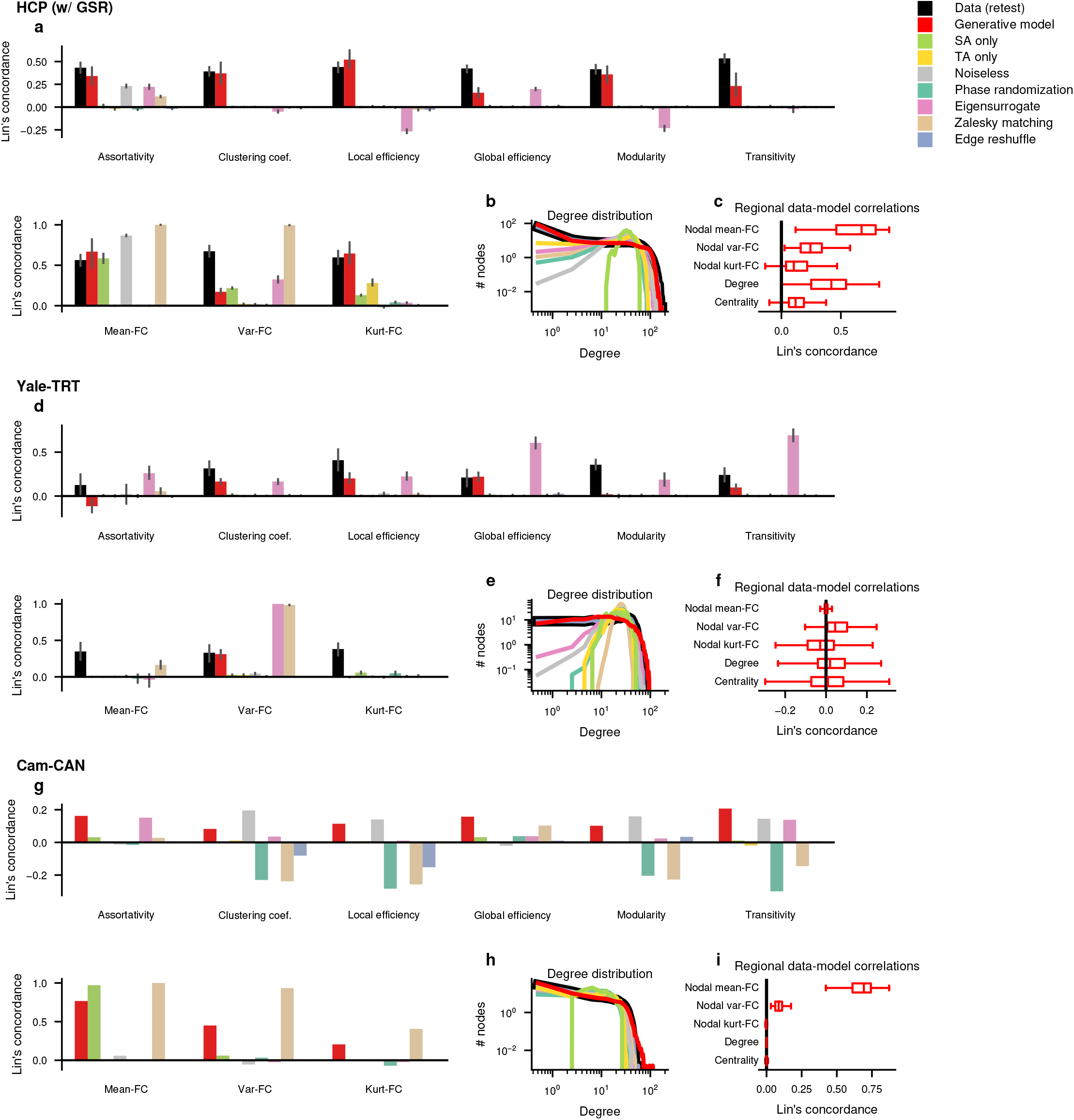
Comparison of model fits for all datasets. For each dataset, the correlation between model and data is shown for timeseries, weighted, and unweighted graph metrics (a,d,g), degree distribution (b,e,h), and distribution of nodal metrics (c,f,i). Statistics of these fits can be found in Table S1.

**Figure S8.**
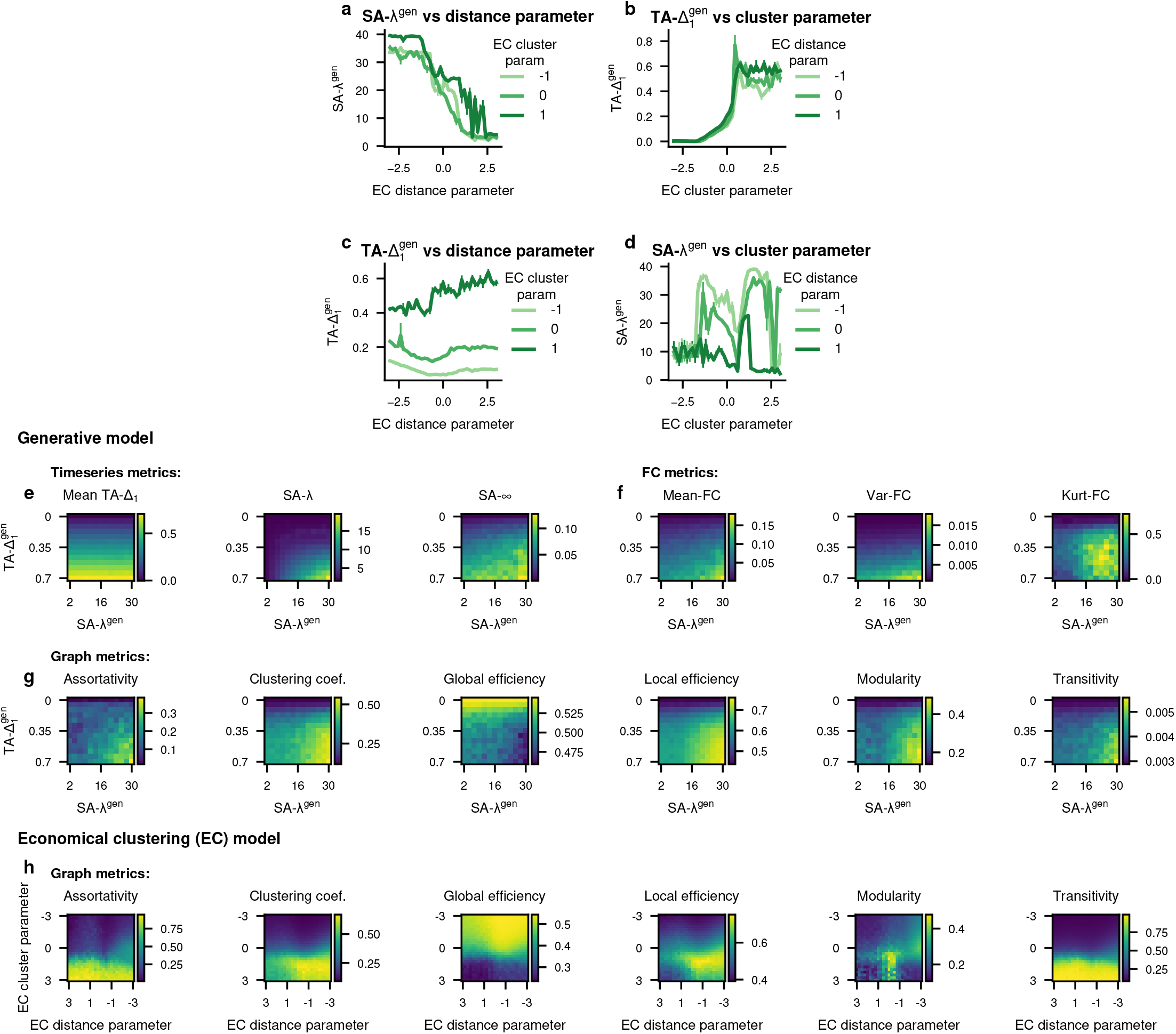
Relationship between the spatiotemporal model and economical clustering (EC) model. (a) For three given values of the EC cluster parameter (γ), the distance parameter (η) was varied and the 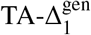 parameter of the best fit spatiotemporal model is shown. (b) For three given values of the EC distance parameter, the EC cluster parameter was varied and the SA-λ^gen^ parameter of the best fit spatiotemporal model is shown. Error bars show standard error across 10 simulations of the EC model. (e-g) The Homogeneous variant of the model was simulated for different values of parameters SA-λ^gen^ and 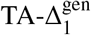. For each timeseries metric (e), weighted graph metric (f), and unweighted graph metric (g), the metric value is plotted as a heatmap. (h) The economical clustering (EC) model was simulated for different values of its distance and cluster parameters. The value of each of the graph metrics is plotted as a heatmap.

**Figure S9.**
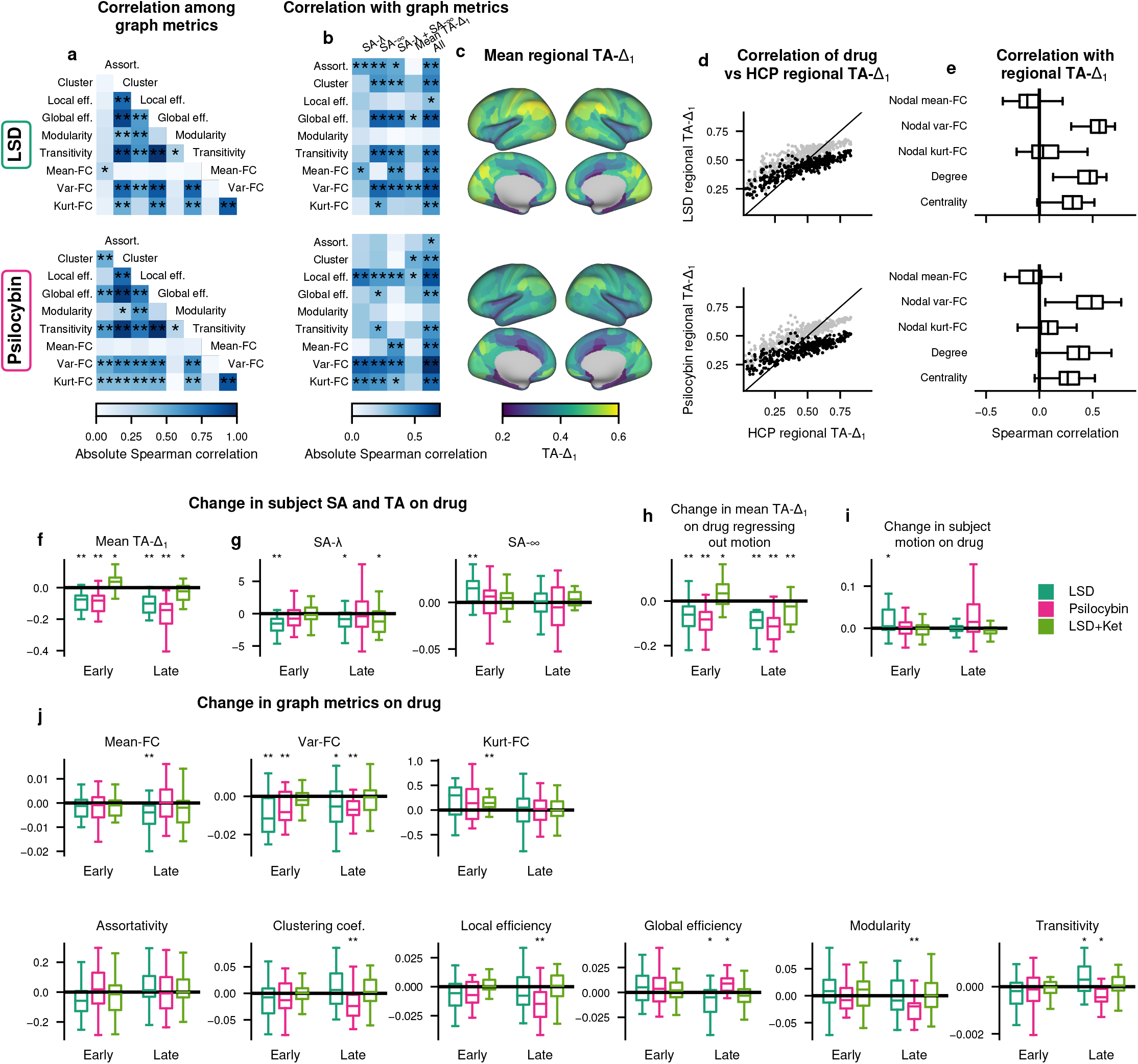
SA and TA under serotonergic modulation. (a-c,e) Similar to Figure 1 and Figure S1, but using only scans performed after administration of LSD (top) or psilocybin (bottom). Due to limited sample size, “SA-λ+SA-∞” and “all” columns in (b) are predicted using leave-one-out cross-validation instead of a training/test split. (d) Correlation of regional TA-Δ_1_ in HCP with those under LSD (top) or psilocybin (bottom). Spearman correlation >.85, significance p<.0001 of the correlation of HCP and LSD or psilocybin regional TA-Δ_1_ determined with SA-preserving scrambles ^8^. Gray points indicate the placebo condition. (f-j) Difference between drug and control across subjects for LSD (blue), psilocybin (green), and LSD with ketanserin (orange), for both early and late scans. Metrics plotted are TA as quantified by TA-Δ_1_ (f), SA as quntified by SA-λ and SA-∞ (g), the residual of a regression model using motion to predict TA-Δ_1_, subject motion as quantified by mean framewise displacement (i), and graph metrics (j). * indicates p<.05, ** indicates p<.01.

**Table S1.**
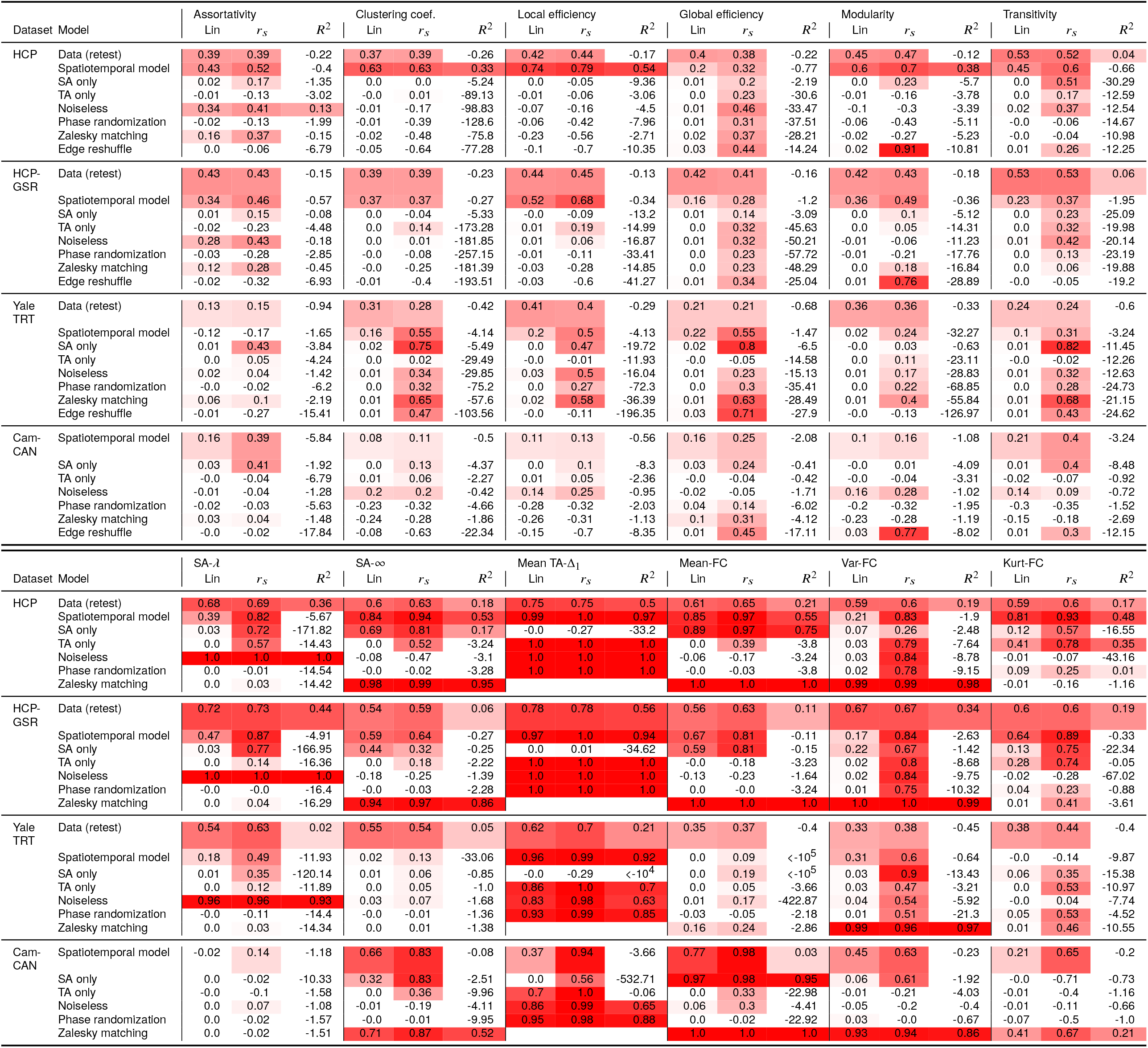
Graph metric fits for all models. For each model, in each dataset, the single subject fits are shown for assortativity, clustering, local efficiency, global efficiency, modularity, and transitivity (top) and SA-λ, SA-∞, TA-Δ_1_, mean-FC, var-FC, and kurt-FC (bottom). Fits are quantified using Lin’s concordance, Spearman correlation *r*_*s*_, and coefficient of determination *R*^2^.

## Footnotes

1 *d* is closely related to other measures of long memory dynamics such as the Hurst exponent *H*; under specific conditions, *H* = *d* +0.5^107^.

